# Propionyl-CoA carboxylase subunit B regulates anti-tumor T cells in a pancreatic cancer mouse model

**DOI:** 10.1101/2023.07.24.550301

**Authors:** Han V. Han, Richard Efem, Barbara Rosati, Kevin Lu, Sara Maimouni, Ya-Ping Jiang, Valeria Montoya, Adrianus W. M. Van Der Velden, Wei-Xing Zong, Richard Z. Lin

**Affiliations:** Department of Physiology and Biophysics, Stony Brook University, Stony Brook, New York; Department of Biomedical Engineering, Stony Brook University, Stony Brook, New York; Northport Veteran Affair Medical Center, Northport, New York, USA; Department of Chemical Biology, Ernest Mario School of Pharmacy, Rutgers-The State University of New Jersey, Piscataway, New Jersey; Department of Microbiology and Immunology, Renaissance School of Medicine at Stony Brook University, Stony Brook, New York, USA; Center for Infectious Diseases, Renaissance School of Medicine at Stony Brook University, Stony Brook, New York, USA

## Abstract

Most human pancreatic ductal adenocarcinoma (PDAC) are not infiltrated with cytotoxic T cells and are highly resistant to immunotherapy. Over 90% of PDAC have oncogenic KRAS mutations, and phosphoinositide 3-kinases (PI3Ks) are direct effectors of KRAS. Our previous study demonstrated that ablation of *Pik3ca* in KPC (*Kras*^G12D^; *Trp53*^R172H^; *Pdx1-Cre*) pancreatic cancer cells induced host T cells to infiltrate and completely eliminate the tumors in a syngeneic orthotopic implantation mouse model. Now, we show that implantation of *Pik3ca*^-/-^ KPC (named αKO) cancer cells induces clonal expansion of cytotoxic T cells infiltrating the pancreatic tumors. To identify potential molecules that can regulate the activity of these anti-tumor T cells, we conducted an *in vivo* genome-wide gene-deletion screen using αKO cells implanted in the mouse pancreas. The result shows that deletion of propionyl-CoA carboxylase subunit B gene (*Pccb*) in αKO cells (named p-αKO) leads to immune evasion, tumor progression and death of host mice. Surprisingly, p-αKO tumors are still infiltrated with clonally expanded CD8^+^ T cells but they are inactive against tumor cells. However, blockade of PD-L1/PD1 interaction reactivated these clonally expanded T cells infiltrating p-αKO tumors, leading to slower tumor progression and improve survival of host mice. These results indicate that *Pccb* can modulate the activity of cytotoxic T cells infiltrating some pancreatic cancers and this understanding may lead to improvement in immunotherapy for this difficult-to-treat cancer.

## INTRODUCTION

Pancreatic ductal adenocarcinoma (PDAC) is one of the most aggressive and lethal cancers, and it is expected to be the 2^nd^ leading cause of cancer death by 2030, with a five-year survival rate of 12% (1). Most PDACs are classified as “cold” tumors without infiltrating T cells and are resistant to currently available immunotherapy with checkpoint inhibitors (2–4). To convert PDAC into “hot” tumors, researchers have explored multiple strategies to promote immune responsiveness of PDAC (5–7). These approaches encompass targeting immune checkpoint inhibitors, including PD1, CTLA4, LAG3, and TIGIT (2,8–11), suppressing immunosuppressive cells such as myeloid-derived suppressor cells (MDSCs) and/or tumor-associated macrophages (TAMs) (12–15), enhancing antigen processing and epitope presentation by promoting MHC class I/II cell surface expression (16–18) and reinstating the expansion and functionality of dendritic cells (19), targeting key factors implicated in stromal fibrosis to facilitate the infiltration of T cells (20–22), and depleting immunosuppressive signals (cytokines and chemokines) in the tumor microenvironment (TME) that hinder the entry and function of T cells (23–26). However, a previous study showed that a minority of PDAC patients have tumors that already exhibit higher levels of infiltrating cytotoxic T cells and this observation is associated with longer survival of these patients (27).

One of the most commonly dysregulated pathways in PDAC is oncogenic KRAS and G12D is the most common KRAS mutation (46%) in PDAC (7,28). Phosphatidylinositol 3-kinases (PI3Ks) are direct effectors of KRAS. There are four PI3K catalytic isoforms and our group have previously shown that PIK3CA isoform plays a critical role in the initiation of PDAC (29) and mediates immune evasion once the tumors are formed (16). Our group reported that KRAS signaling through PIK3CA can reduce the expression of MHC class I molecules, thus reducing antigen recognition and T cell infiltration (16). We showed that genetic ablation of PIK3CA in KPC (*Kras^G12D^;Trp53^R172H^;Pdx1-Cre*) pancreatic tumor cells led to T cell recognition and complete elimination by the host immune system in a syngeneic orthotopic implantation mouse model (16). Using this well-characterized syngeneic orthotopic implantation mouse model with the αKO (*Kras^G12D^;Trp53^R172H^;Pdx1-Cre;Pik3ca^-/-^*) cell line and single-cell sequencing analysis, we found clonal expansion of cytotoxic T cells infiltrating αKO tumors. To investigate potential molecules within αKO cells that affect the activity of host T cells, we performed an *in vivo* genome-wide CRISPR gene-deletion screen using αKO cells and found that deletion of propionyl-CoA carboxylase subunit B (PCCB) could reverse the immune-elimination phenotype. We further discovered that T cells infiltrating the p-αKO (*Kras^G12D^;Trp53^R172H^;Pdx1-Cre;Pik3ca^-/-^;Pccb^-/-^*) tumors were suppressed and could not eliminate the tumor cells. However, these exhausted anti-tumor T cells could be reactivated by neutralizing the PD-L1/PD1 axis, restoring the ability of these immune cells to eliminate the pancreatic tumor.

## RESULTS

### Clonal expansion of cytotoxic T cells infiltrating the pancreatic αKO tumors

In our previous study, we demonstrated that αKO tumors, when implanted in immunocompetent C57BL/6 (B6) mice, are recognized and eliminated by the host immune system, with cytotoxic T cells playing a pivotal role in this process (16). To investigate the status of the cytotoxic T cells infiltrating to the tumor microenvironment (TME), single cell RNA sequencing (scRNA-seq) was conducted in combination with concurrent T cell receptor (TCR) repertoire sequencing. On day #12 post implantation in two B6 mice, both of the αKO tumors were harvested and dissociated to generate single cell suspensions. CD8^+^ T cells were isolated using microbead preparations, and both the transcriptional expression and TCR repertoire profiles were obtained using 10x genomic protocols (Figure 1A). Following stringent quality control filters (outlined in the Methods section), we successfully obtained gene expression profiles and TCR sequences from the same cytotoxic T cells.

**Figure 1.**
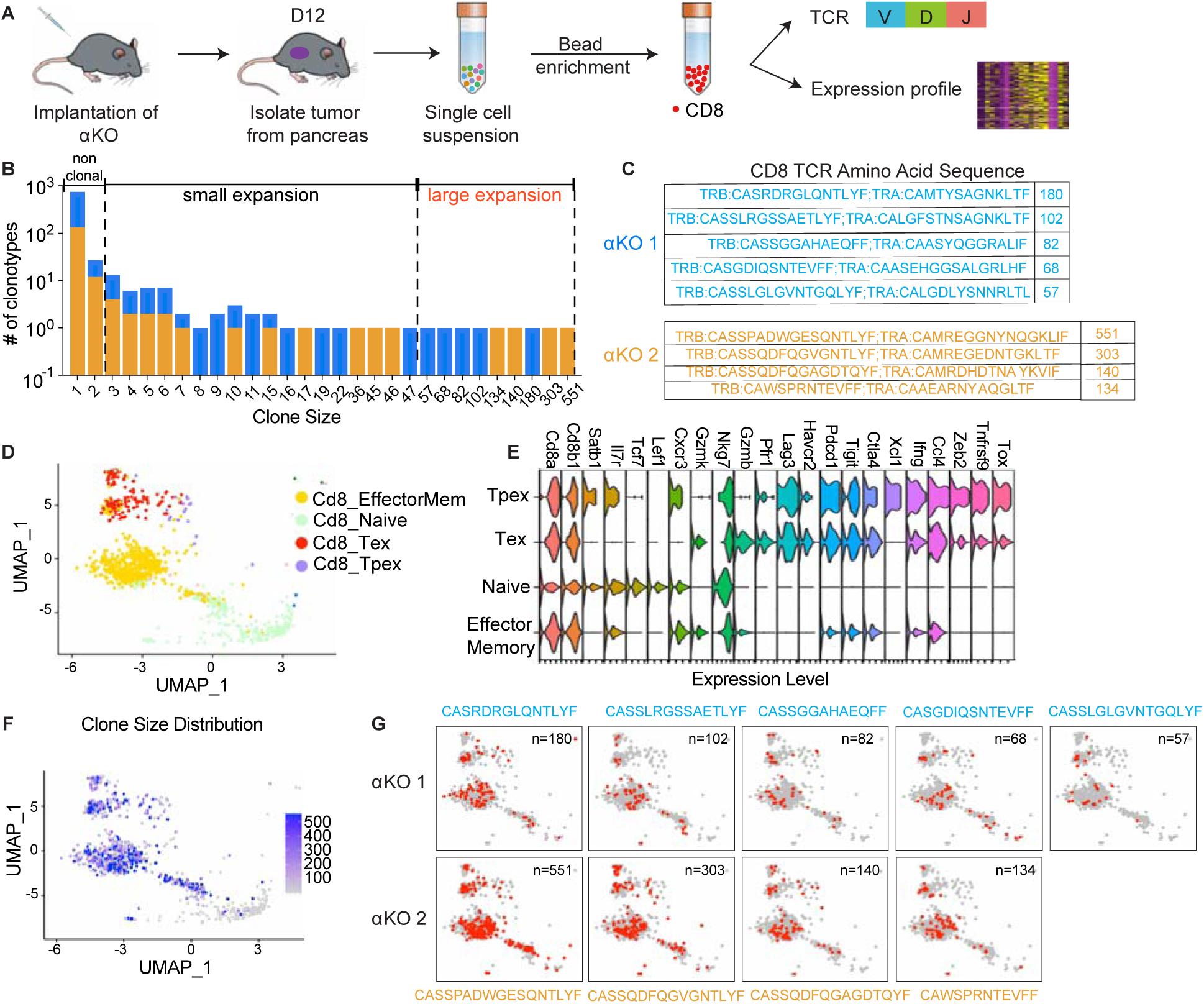
Clonal expansions of cytotoxic T cells infiltrating the αKO pancreatic tumors. **(A)** Experimental design schema for single cell RNA sequencing of infiltrating CD8^+^ T cells in αKO tumors. Cells were implanted in the pancreas of two B6 mice and tumors were harvested at 12 days post-implantation. Single cell suspensions were enriched with CD8^+^ beads followed by concurrent single-cell transcriptional and T-cell receptor (TCR) profiling using 10x Genomics protocols. **(B)** Size distribution for CD8^+^ T cell clonal types in αKO tumors. Non-clonal: N≤2; small expansion: 3 ≤N<50; large expansion: N≥50. **(C)** Amino acid sequences of the CDR3 region for large expansion clonotypes. αKO1 tumor has 5 and αKO2 tumor has 4 large expansion clonotypes (full list shown in the Supplementary File 1). **(D)** UMAP of all CD8^+^ T cell clonotypes. Principle component analysis of gene expression profiles generated 4 cell clusters annotated based by ProjectTILs. EffectorMem: effector memory; Tex: exhausted; Tpex: precursor exhausted; NaiveLike: naïve. Each dot corresponds to a single cell. **(E)** Major markers used for cell type annotation by Projectils. **(F)** Clonal size distribution mapped to the gene expression UMAP showing that the largest clonotypes are mostly effector memory T cells. **(G)** The gene expression profile distribution of individual large expansion clonotypes are shown. The majority of cells for each clone mapped to the effector memory T cell functional group.

The diverse expression of TCR repertories is crucial for effective adaptive immune responses. Sequencing the TCR repertoire allows us to identify clonotypic diversity and detect clonal expansion (30). Analysis of the CD8^+^ T cells’ clonal size distribution revealed clonal expansions in both αKO tumors, suggesting T cells in both tumors have been activated in response to tumor antigen (Figure 1B). A clonotype is defined as 3 or more cells with identical TCR sequences. A clonal size of 3 to 49 is defined as small expansion, and a clonal size of 50 or more is designated as large expansion (Figure 1B). Figure 1C displays the TCR sequences for CD8^+^ clones with large expansions (clonal size ≥50) in both tumors.

We next analyzed the clonality of these cells with respect to their transcriptomes. The gene expression profiles were visualized using the uniform manifold approximation and projection (UMAP) (Figure 1C). To achieve this, the expression data underwent unsupervised clustering by Seurat. Cell type annotations were determined using ProjectTILs, a computational method that annotates cell types by projecting scRNA-seq data onto reference profiles constructed from well-established canonical cell markers and clinical data (31). The Violin Plot in Figure 1E illustrates major markers used for cell type annotation by ProjectTILs. Based on the annotation, clusters are annotated as effector memory CD8^+^ T cells (Cd8_EffectorMem), naïve-like CD8^+^ T cells (Cd8_NaiveLike), exhausted CD8^+^ T cells (Cd8_Tex), and precursor exhausted CD8^+^ T cells (Cd8_Tpex). We next mapped the TCR profiling datasets to the single-cell gene expression datasets and then color coded by the size of each clonotype (Figure 1F). The degree of expansion observed among the clonotypes associated strongly with the phenotypic clusters of T cells, as large clones are mapped to the Cd8_EffectorMem and Cd8_Tex, while nonclonal T cells mapped to the Naïve T cell group (Figure 1F). Figure 1G shows the single-cell gene profile distribution of large expansion (clonal size ≥50) clonotype from both tumors. Taken together, these results show that there is a massive expansion of Cd8_EffectorMem clones and these activated anti-tumor T cells are likely are responsible for the immune-mediated elimination of αKO pancreatic tumors.

### A genome-wide CRISPR gene-deletion screen to identify molecules contributing to Pik3ca-mediated pancreatic tumor immune evasion

To identify potential molecules in tumor cells that can regulate the activity of these anti-tumor clonal T cells, we conducted an *in vivo* genome-wide gene-deletion screen using αKO cells implanted in the mouse pancreas. A schema for the CRISPR knockout screen is shown in Figure 2A. Briefly, αKO cells were first infected with the Genome-Scale CRISPR Knockout (GeCKO v2) pooled lentiviral library A, which targets the complete mouse genome of 20,611 genes with three sgRNAs for each gene, as well as four sgRNAs for each of the 1175 miRNAs and 1000 control sgRNAs. Equal portion of the mixed cell pool was implanted into the pancreas of eight immunocompetent B6 mice and tumor progression was monitored longitudinally by an IVIS Lumina III *in vivo* imaging system (Figure 2B). When implanted in B6 mice, αKO tumors completely regressed. Therefore, cells with genetically deleted molecules that can reverse the immune recognition phenotype should lead to tumor progression. Indeed, 2 out of 8 mice eight weeks after cell implantation exhibited tumor progression, which were labeled as tumor #1 and tumor #2 (Figure 2B). Tumors were harvested from both mice for further analysis. The remaining mice (6 out 8) exhibited complete tumor regression consistent with the phenotype we previously observed with αKO cells (Figure 2B). Genomic DNA was prepared from both tumors and subjected to next generation sequencing (NGS) to detect the presence of incorporated sgRNA sequences. Sequenced data was analyzed using Model-based Analysis of Genome-wide CRISPR/Cas9 Knockout (MAGeCK), and the counts for each sgRNA sequence are shown in Figure 2C (full list shown in the Supplementary File 2). Somewhat surprisingly, four common sgRNA sequences were identified in both tumors. The four targeted genes are *Pccb* (propionyl-CoA carboxylase), *Nr6a1* (nuclear receptor subfamily 6 group A member 1), *Ldoc1l* (leucine-zipper protein 1), and *Adck3* (aarF-domain-containing kinase 3). Tumor #2 has two additional sgRNA sequences targeting two other genes that were not present in Tumor #1. Despite the fact that tumor #1 and tumor #2 grew at different rates and were harvested at different time points, week 7 and week 13 respectively, the detected sgRNAs were highly similar, strongly suggesting that the identified genes play critical roles in *Pik3ca*-mediated immune evasion.

**Figure 2.**
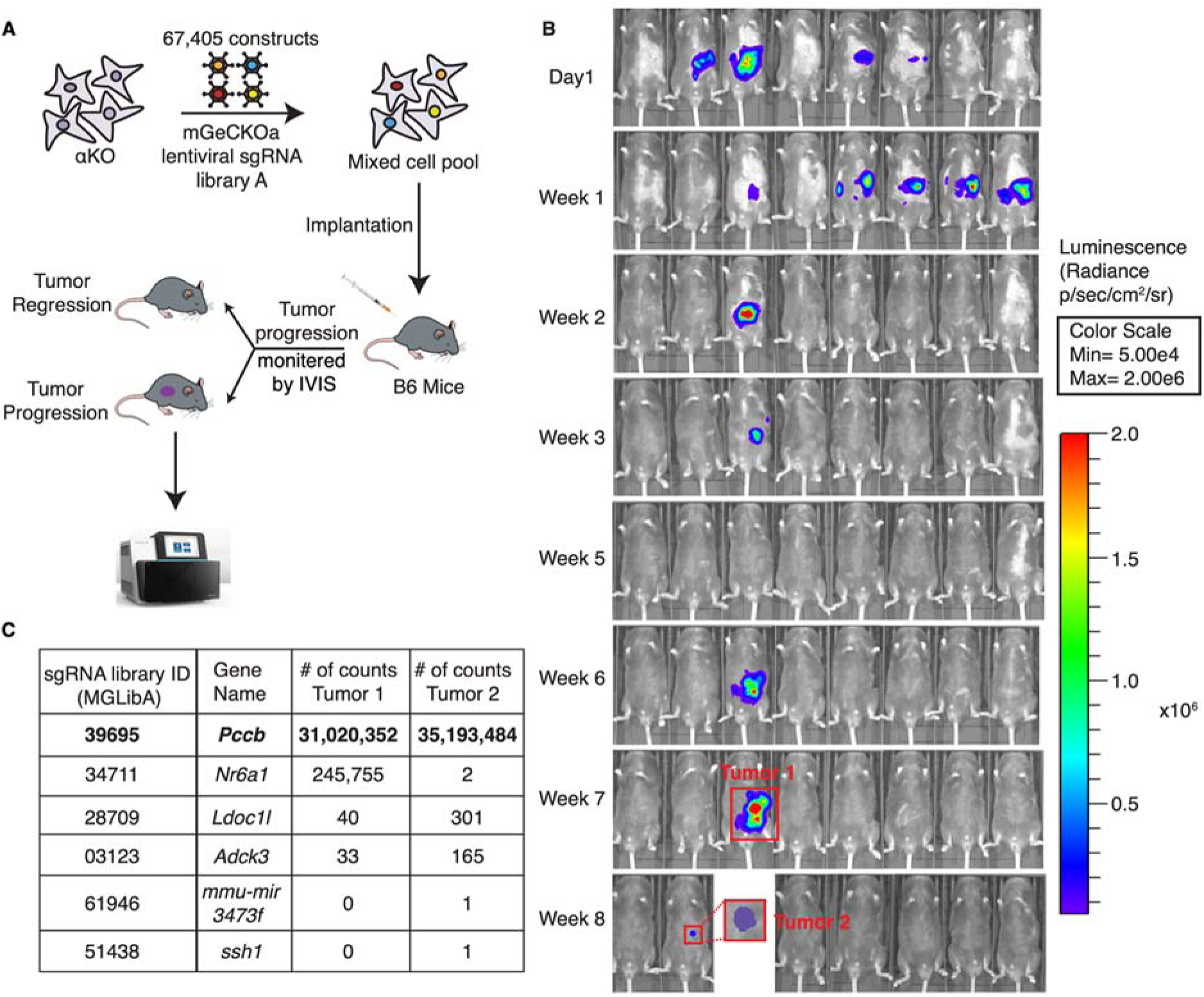
A genome-wide CRISPR gene-deletion screen identifies *Pccb* contributing to PIK3CA-mediated pancreatic tumor immune evasion. (A) A schema for the genetic screen. Mouse genome-scale CRISPR/Cas9 knockout library (mGeCKO v2A) containing three sgRNA each for 20611 genes was transduced into *Kras^G12D/+^;Tp53^R172H/+^;Pik3ca^-/-^* (αKO) cells. Pooled cells were implanted in the pancreas of eight C57BL/6J mice. **(B)** IVIS imaging was performed to monitor tumor progression in these mice. In two animals, tumors were still present at the end of the screen (annotated as tumor 1 & 2). **(C)** Harvested DNAs prepared from the tumors were subjected to next generation sequencing (NGS) to identify enriched sgRNAs barcodes. Sequences corresponding to only six genes in the mGeCKO v2A library have detectable NGS counts as shown in the table. In both tumors, sgRNA sequences mapping to the *Pccb* gene are most numerous (full list shown in the Supplementary File 2).

In both tumors, *Pccb* sgRNA sequence MGLibA ID 39695 was detected at many orders of magnitude higher than other three sgRNA sequences (Figure 2C). Propionyl-CoA carboxylase (PCC) has two subunits PCCA (propionyl-CoA carboxylase subunit A) and PCCB and catalyzes the carboxylation of propionyl-CoA to methylmalonyl-CoA, which is then converted to succinyl-CoA that enters the tricarboxylic acid cycle (TCA). PCC dysfunction leads to a rare genetic disease called propionic acidemia, which has been extensively studied (32–34). However, PCCB’s role in cancer and whether it plays a role in the tumor immune response is completely unknown.

### Ablation of PCCB reverses the immune recognition of αKO cells

To validate the genetic screen result, we next genetically ablated *Pccb* in αKO cells using CRISPR/Cas9 techniques and generated the PCCB-null p-αKO cell line. PCCB ablation was confirmed by western blotting (Figure 3A). We then orthotopically implanted p-αKO cells in the head of pancreas of B6 mice and monitored tumor growth by IVIS imaging (Figure 3B). Most of the mice exhibited tumor formation within one week. In some of the host animals, tumor regressed when they were reimaged at week #2. However, in all of the animals, pancreatic tumors eventually progressed and 9 out of 12 mice died by week #8 when the experiment was stopped (Figure 3B & C). Postmortem examination showed that the three surviving mice had large pancreatic tumors that were also detectable by IVIS imaging (Figure 3B). In comparison, while 100% of mice implanted with KPC cells died by week #3, all mice implanted with αKO cells remained tumor free by week #8 (Figure 3C & D). These results indicate that PCCB loss can reverse immune recognition of αKO cells.

**Figure 3.**
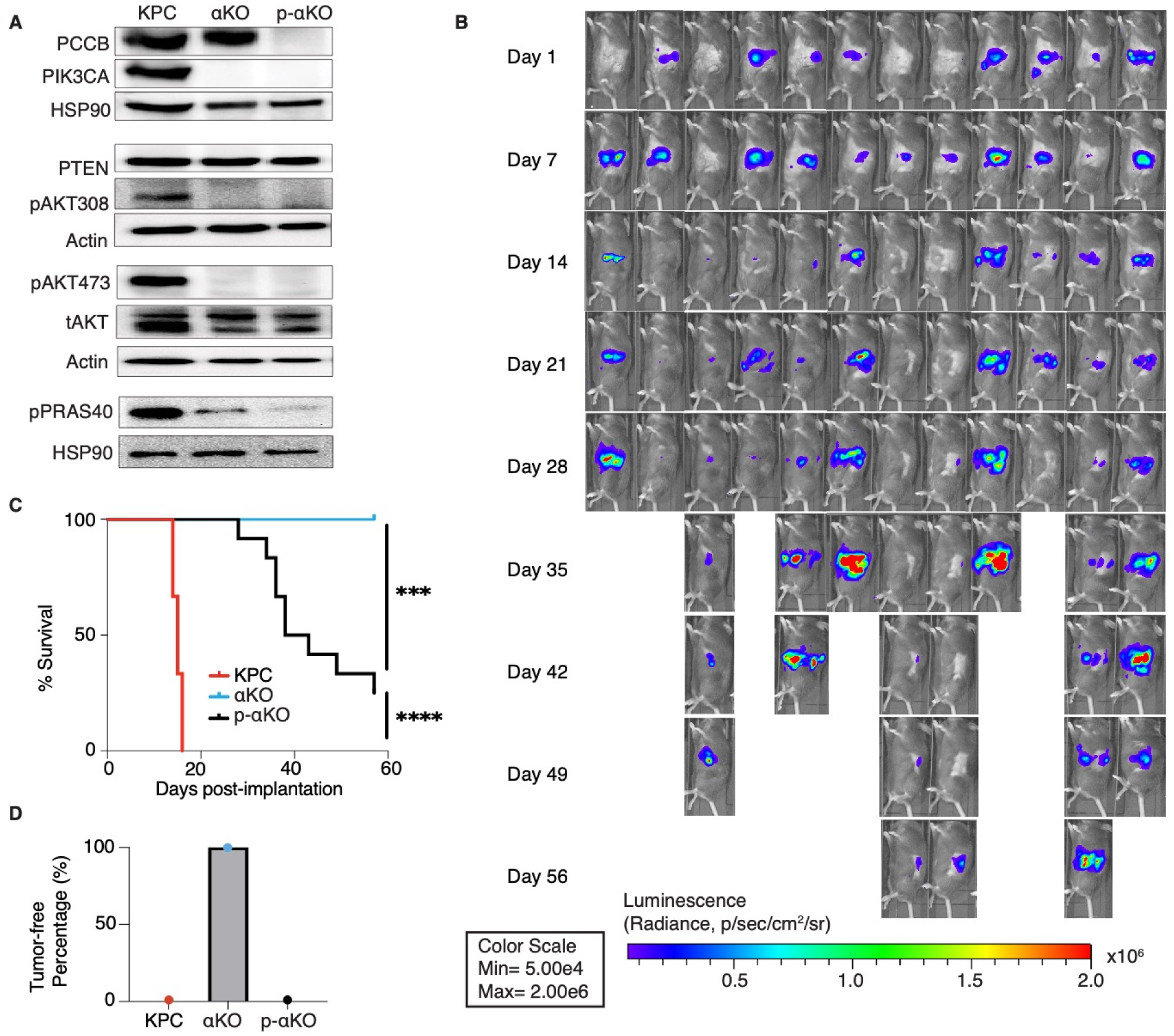
Ablation of PCCB reverses the immune recognition of αKO cells. **(A)** *Pccb* was deleted in αKO cells by CRISPR/Cas9 to generate p-αKO cells. Western blotting confirmed PCCB and PIK3CA loss in p-αKO cells. Additionally, western blotting was performed for PTEN, phospho-Thr308 AKT, phospho-Ser473 AKT, and phospho-PRAS40, with Actin and HSP90 used as loading controls. **(B)** Orthotopic implantation of p-αKO tumors in the pancreas of C57BL/6J mice was monitored by IVIS imaging of the luciferase signal weekly for 8 weeks. **(C)** Kaplan-Meier survival curves for mice implanted with KPC, αKO, and p-αKO cells. Median survival KPC: 15.5 days (N=8); αKO, all alive (N=9); p-αKO, 40.5 days (N=12). ****P < 0.0001 for KPC vs. αKO; ***P=0.0009 for αKO vs. p-αKO; ****P < 0.0001 for KPC vs. p-αKO (log-rank test). **(D)** Percentage of B6 mice that are tumor-free at 8 weeks after implantation with KPC (0%), αKO (100%) and p-αKO (0%) cells.

In standard 2D culture, p-αKO cells grew at a similar rate compared to αKO cells (Figure 3 – figure supplement 1A). Note that both αKO and p-αKO cell lines exhibited significantly slower growth rate than KPC cells. This reduced growth rate explains, at least in part, the longer survival time of p-αKO implanted mice compared with KPC-implanted mice, where the median survival of KPC was 15.5 days and the median survival of p-αKO 40.5 days (Figure 3C). Immunoblotting showed no changes in phosphorylation of PIK3CA’s downstream effector AKT at both Thr308 and Ser473 residues. As expected, phosphorylation of PRAS40, which is a direct effector of AKT, was also downregulated in both cell lines lacking PIK3CA as compared to KPC cells. PTEN is a negative regulator of the PI3K-AKT pathway, and its expression was not different between the three cell lines (Figure 3B). In summary, the loss of PCCB did not reverse the changes in PIK3CA-AKT signaling in αKO cells. These results indicate that progression of p-αKO tumors is through a pathway independent of PI3K and AKT. In addition, the mitochondrial function evaluated by measuring the oxygen consumption rate (OCR) with Seahorse, as well as the intracellular glycolysis and TCA metabolite changes measured by untargeted liquid chromatography-mass spectrometry (LC/MS) both showed comparable levels between αKO and p-αKO cell lines, indicating that the increased growth of p-αKO cells *in vivo* is probably not due to PCCB-related mitochondrial metabolic changes (Figure 3 – figure supplement 1B & E).

### T cells infiltrate p-αKO tumors with increased expression of immune checkpoints

Our previous study demonstrated that regression of αKO tumors is due to T cell infiltration and subsequent elimination by the host immune system (16). We performed immunohistochemistry (IHC) to assess the status of infiltrating T cells in p-αKO, αKO, and KPC tumors. After implantation, IVIS imaging was used to confirm presence of tumors, which were then harvested for analysis. H&E staining confirmed tumor formation (Figure 4A) and immediate adjacent tissue sections were stained for infiltrating T cells using anti-CD3, CD4 and CD8 antibodies. Unexpectedly, p-αKO tumors were also highly infiltrated with T cells, similar to αKO tumors, and as expected, KPC tumors were mostly devoid of T cells (Figure 4A & B). Therefore, the conversion of cold KPC tumor into inflamed αKO tumor upon genetic deletion of *Pik3ca* in KPC was not reversed by ablation of PCCB.

**Figure 4.**
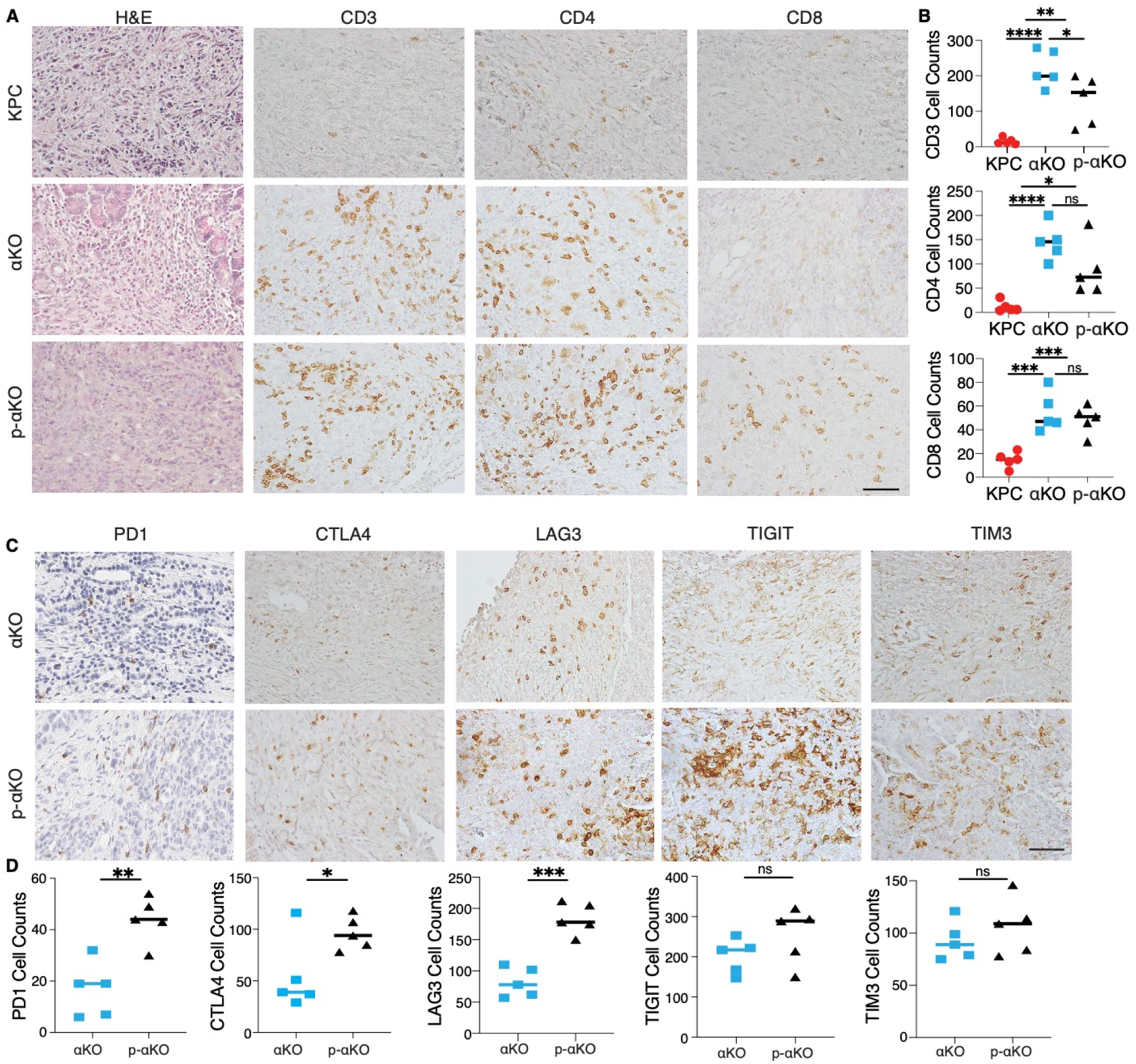
T cells infiltrate p-αKO tumors with increased expression of immune checkpoints. **(A)** Pancreatic tissue sections prepared from C57BL/6J mice implanted with KPC, αKO, and p-αKO cells. Tumor sections were stained with H&E and IHC with CD3, CD4, and CD8 antibodies. Representative sections are shown. Scale bar: 60μm. **(B)** Quantification of tumor-infiltrating T cells in 5 representative tumor sections for each group (mean ± SD, n=3). CD3: ****P<0.0001 for KPC vs. αKO; **P=0.0063 for KPC vs. p-αKO; *P=0.0471 for αKO vs. p-αKO. CD4: ****P<0.0001 for KPC vs. αKO; *P=0.0164 for KPC vs. p-αKO; ns for αKO vs. p-αKO. CD8: ***P=0.0009 for KPC vs. αKO; ***P=0.0005 for KPC vs. p-αKO; and ns for αKO vs. p-αKO (two-tailed t test). ns: not significant. **(C)** αKO and p-αKO cells were implanted in the head of pancreas of C57BL/6J mice. Tumor sections were stained with checkpoint markers: PD1, CTLA4, LAG3, TIGIT and TIM3. Representative sections are shown. Scale bars: 60 μm. **(D)** Quantifications of cells positive for each checkpoint marker at representative tumor sections (mean ± SD, N=5). αKO vs. p-αKO: **P = 0.0023 for PD1; *P = 0.0371 for CTLA4; ***P = 0.0002 for LAG3; ns for TIGIT and TIM3.

As high expression of checkpoint inhibitory receptors is associated with reduced ability to perform cytotoxic functions of CD8^+^ T cells, we examined whether T cell checkpoint inhibitory receptors, including programmed cell death 1 (PD-1), cytotoxic T-lymphocyte associated protein 4 (CTLA-4), T cell immunoreceptor with Ig and ITIM domains (TIGIT), lymphocyte activating 3 (LAG3), and hepatitis A virus cellular receptor 2 (TIM3) were altered upon PCCB ablation. Tumor tissues harvested from mice implanted with αKO and p-αKO were analyzed by IHC cells positive for the stained markers were quantified (Figure 4C). In contrast to αKO tumor sections, p-αKO tumors displayed markedly elevated cell counts positive for PD1, CTLA4 and LAG3. However, despite comparable cell counts, the expression levels of TIGIT and TIM3 appeared stronger in p-αKO tumor sections (Figure 4D). These results suggest that infiltrating CD8^+^ T cells in p-αKO tumors might be functionally exhausted with compromised anti-tumor cytotoxic activity.

To explore the potential involvement of other immunosuppressive cells, flow cytometry analysis was conducted on myeloid cells isolated from αKO and p-αKO tumors. The results revealed a significant twofold increase in M2 macrophage infiltration within the p-αKO tumors. This finding provides additional support for the presence of an immunosuppressive TME specifically within the p-αKO tumors, indicating the complexity of the immune landscape in pancreatic cancer (Figure 4 – figure supplement 1A & B).

### Inhibition of PD1/PD-L1 checkpoint leads to elimination of most p-αKO tumors

Although multiple immune checkpoints are upregulated in p-αKO tumors, we investigated whether blocking the PD1/PD-L1 immune checkpoint alone would reactivate some of the anti-tumor T cells. We implanted p-αKO cells into a PD1-null mouse line bred in the B6 genetic background (Figure 5A). After 8 weeks when the experiment ended, 8 out 9 mice were still alive and 5 of the animals were tumor free (Figure 5B & C). We next tested if pharmacological blockade of PD1/PD-L1 would achieve a similar result. We implanted p-αKO cells in wild-type B6 mice and then treated them weekly with a neutralizing anti-PD-1 antibody or vehicle control (Figure 5A). After 8 weeks of treatment, 9 out 13 anti-PD1-treated mice were alive whereas only 2 out 9 vehicle-treated mice survived (Figure 5B). Four mice in the anti-PD1-treated group remained tumor free. In contrast, all vehicle-treated mice developed large pancreatic tumors including the two surviving mice at week #8 (Figure 5C).

**Figure 5.**
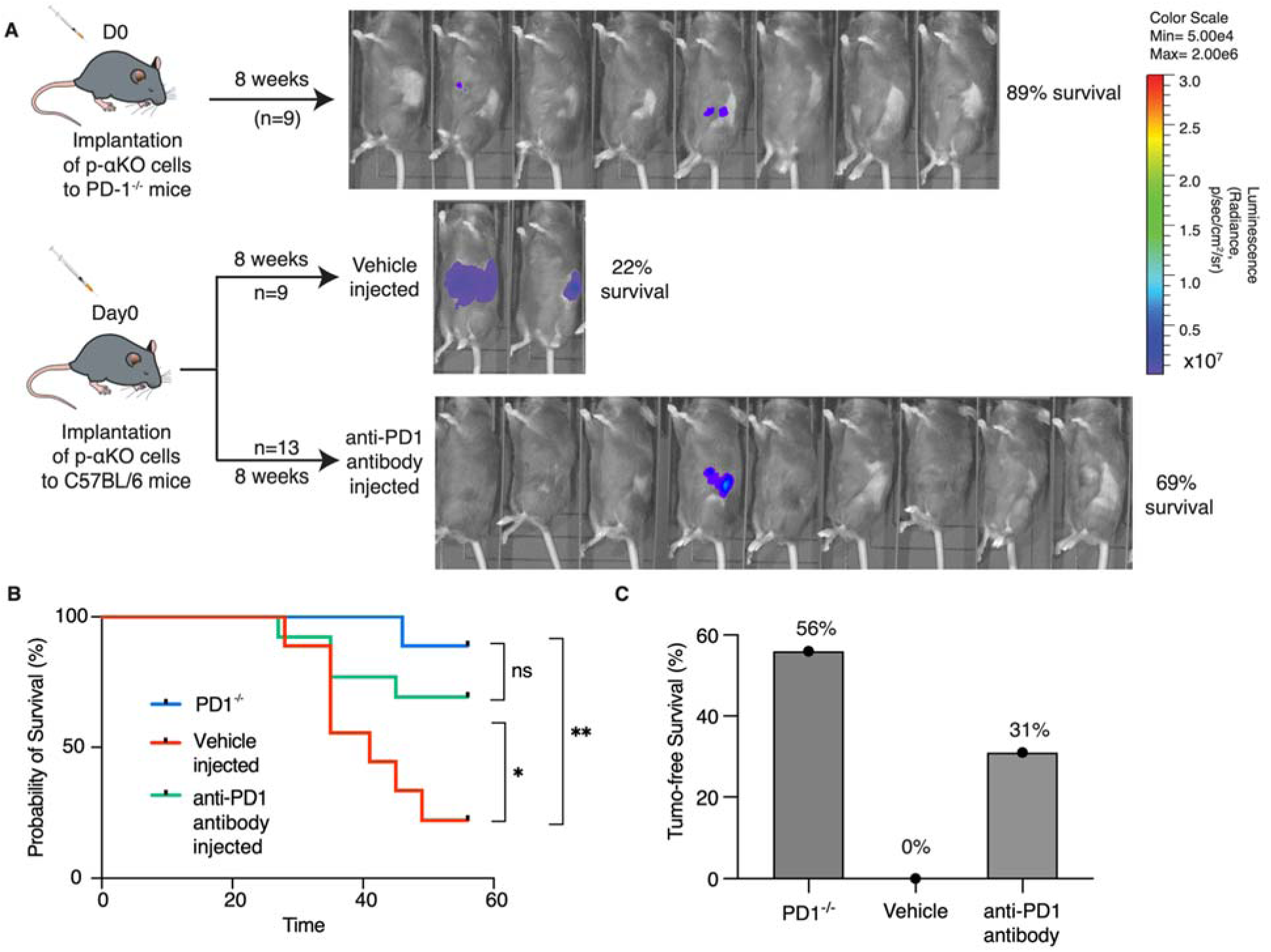
Inhibition of PD-1/PD-L1 checkpoint interaction leads to elimination of most p-αKO tumors. **(A)** PD1^-/-^ mice (N = 9) were implanted with p-αKO cells in the pancreas and was monitored for 8 weeks. IVIS imaging was performed to assess tumor size at the end of the experiment. C57BL/6J mice (N = 22) were implanted with p-αKO cells in the pancreas and randomly divided into two groups. One group was injected with 12.5 mg/mg of PD1 neutralizing antibody weekly for 8 weeks, and the vehicle group was injected with the inVivo Pure dilution buffer. IVIS imaging was performed to assess tumor size at the end of the experiment. **(B)** Kaplan-Meier survival curves for PD1^-/-^ mice implanted with p-αKO cells, and vehicle and anti-PD1 treated C57BL/6J mice implanted with p-αKO cells. Survival rate for PD1^-/-^ group is 89% and median survival undefined; survival rate for vehicle injected group is 22% and median survival is 41 days; survival rate for anti-PD1 injected group is 69% and median survival undefined. *P = 0.0436 for vehicle vs. anti-PD1 antibody injection. **(C)** Tumor free survival percentage of PD1^-/-^ (56%), vehicle injected (0%), and anti-PD1 antibody injected (31%) groups.

### Simultaneous analysis of anti-tumor cytotoxic T cell infiltrating p-αKO tumors by scRNAseq and TCR repertoire sequencing

To investigate if p-αKO tumors, like αKO cells, also induced clonal expansion of cytotoxic T cells, we performed a scRNA-seq analysis (same protocol as Figure 1) on CD8^+^ T cells isolated from p-αKO tumors implanted in WT mice. In addition, to assess if PD1/PD-L1 blockade reactivated anti-tumor T cells, we also performed the same analysis on CD8^+^ T cells isolated from p-αKO tumors implanted in PD1-null mice (n=2 for each group). Similar to what we observed in αKO tumors (see Figure 1), there are large clonal expansions of CD8^+^ T cells in p-αKO tumors implanted in either WT or PD1-null mice (Figure 6A).

**Figure 6.**
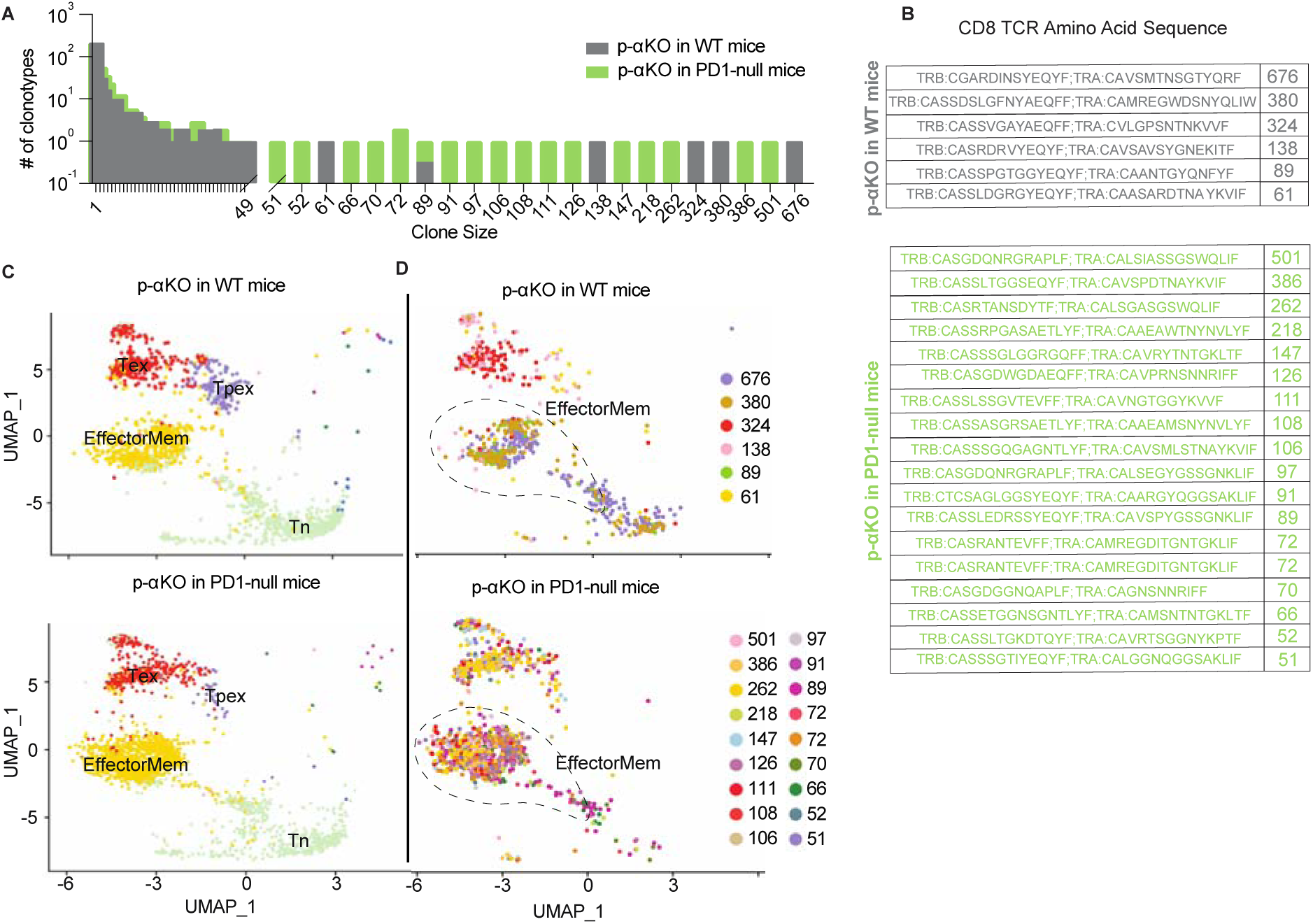
Increased number of anti-tumor cytotoxic T cells infiltrating p-αKO tumors implanted in PD1-null vs. WT mice. WT and PD1-null mice in B6 genetic background were implanted with p-αKO cells in the pancreas (n=2 for each group). Tumors were harvested from these animals 12 days post implantation. T cell suspensions prepared from these tumors were subjected to scRNAseq and TCR sequencing analysis using the same protocol as Figure 1. **(A)** The bar graph shows size distribution of CD8^+^ T cell clonotypes found in WT vs. PD1-null tumors. **(B)** CDR3 sequences of clonotypes with size ≥50 (defined as large expansion). Tumors in WT mice had 6 clonotypes with size ≥50 cells and tumors in PD1-null mice had 18 clonotypes with size ≥50 cells. **(C)** UMAP showing gene expression profiles of all CD8^+^ T clonotypes found in both groups. EffectorMem: effector memory; Tex: exhausted; Tpex: precursor exhausted; Tn: naïve. Each dot corresponds to a single cell. **(D)** UMAP showing gene expression profiles of large expansion CD8^+^ T cell clonotypes. Each clonotype is color coded showing the functional mapping of individual cells. The dash line marks where effector memory T cells are located on the UMAP.

The TCR sequences of the large expansions (≥50 cells per clonotype) in both groups are shown in Figure 6B. Interestingly, the number of clonotypes with large expansions are much higher in tumors implanted in PD1-null mice. Indeed, the number of clonotypes observed in p-αKO tumors implanted in PD1-null mice are even greater than those found in αKO tumors implanted in WT mice.

Transcriptomic data from all clonal CD8^+^ T cells were then clustered in an unsupervised fashion and visualized by UMAP (see methods section) and annotated the clusters by ProjecTILs to identify effector memory T cells (EffectorMem), naïve T cells (Tn), exhausted T cells (Tex) and precursor exhausted T cell (Tpex) groups (Figure 6C). Surprisingly, we observed large number of clonal CD8^+^ EffectorMem and Tex cells in p-αKO tumors implanted in either WT or PD1-null mice. We next mapped clonotypes with large clonal expansions (≥50 cells) to the transcriptomic data and displayed the results by clonotype (Figure 6D). Again, in both groups large expansion clonotypes mainly mapped to EffectorMem and Tex gene expression profiles. There are many more CD8^+^ clonotypes that mapped to the EffectorMem in tumors implanted in PD1-null mice as compared to WT mice. Notably, 856 CD8^+^ T cells (all 18 large expansion clonotypes) were observed in the EffectorMem group for tumors implanted in PD1-null mice, whereas only 361 cells (from 3 clones) were observed appeared in tumors implanted in WT mice (Figure 6E). Indeed, we found more CD8^+^ EffectorMem cells in p-αKO tumors implanted in PD1-null mice then αKO tumors implanted in WT mice (856 vs. 559). Taken together, these results suggest that regression of p-αKO tumors implanted in mice with PD1/PD-L1 blockade is due to reactivation of anti-tumor cytotoxic T cells.

Recent studies propose that immune checkpoint blockade, such as anti-PD1, promotes Tpex to generate effector T cells (35–37). Our findings is consistenet with this hypothesis, p-αKO tumors implanted in WT mice exhibit a distinct population of Tpex. Treatment with an anti-PD1 antibody resulted in their transformation into effector memory T cells, as observed in p-αKO tumors implanted in the PD1-null mice.

### PCCB expression in human pancreatic cancer

To explore the potential significance of *PCCB* expression in PDAC patients, we queried the Cancer Genome Altas Program (TCGA) database and compared the overall survival of patients with high *PCCB/PIK3CA* ratio to patients with low *PCCB/PIK3CA* ratio. Compared with patients with lower *PCCB/PIK3CA* ratio, patients with higher ratio trended to have better survival (Figure 7A). Additional clinical data of PDAC patients were obtained from gene expression omnibus (GEO) database. The GSE15471 is an expression analysis of 36 PDAC patients from the Clinical Institute Funding, where tumors and matching normal pancreatic tissue samples were obtained at the time of surgery. The GSE16515 is the expression data from Mayo Clinic, and it consists of 36 tumor samples and 16 normal samples. Both data revealed significantly lower *PCCB* expressions in tumor sections compared with adjacent normal tissues (Figure 7B), which is consistent with our observation that low *PCCB* levels lead to tumor progression.

**Figure 7.**
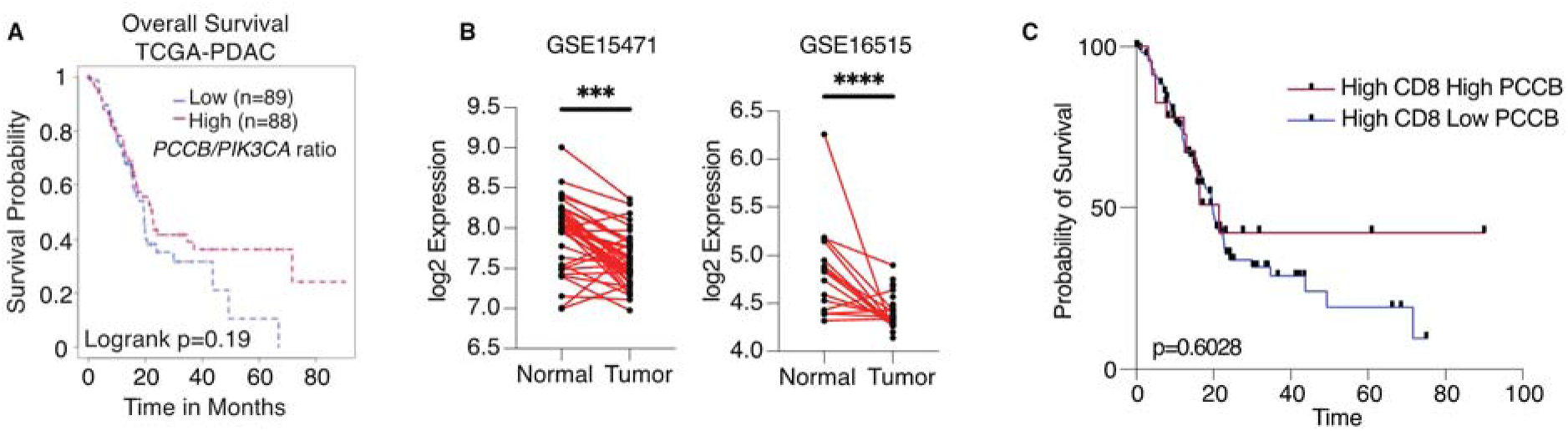
PCCB regulates PDAC survival in a TIL-dependent manner. **(A)** Overall survival between patients with high *PCCB/PIK3CA* ratio vs. patients with low *PCCB/PIK3CA* of 177 PDAC patients from TCGA database. P=0.19. **(D)** Two PDAC clinical data GSE15471 and GSE16515 illustrating *PCCB* mRNA expression in tumor sections vs. adjacent normal tissues. ***P=0.0001 for GSE15471, N=36 for tumor, and N=36 for normal; and ****P<0.0001 for GSE16515, n=36 for tumor, and N=16 for normal. **(C)** The overall survival comparisons among PDAC patients with high infiltrating CD8 T cell level with high or low PCCB expression in TCGA dataset analyzed with TCGA database. High vs. low *PCCB* expression was defined as 80% percentile.

## DISCUSSION

The PI3K signaling pathway plays an important role in regulating the immune microenvironment in PDAC, and previous studies have indicated that specific PIK3CA inhibitors hold promise in transforming immune-deficient cold tumors into inflamed, hot tumors (38–43). Our previous investigations with a PDAC mouse model revealed that downregulation of PIK3CA in tumor cells enhances their recognition and elimination by T cells (16). Building on this foundation, our present study demonstrates that implanting αKO cells into B6 mice induces a clonal expansion of cytotoxic T cells within the TME.

Conducting a genome-wide *in vivo* CRISPR screen with αKO cells implanted in the mouse pancreas allowed us to identify key molecules governing the activity of these anti-tumor T cells. Notably, we discovered that the loss of PCCB, a subunit of the mitochondrial enzyme propionyl-CoA carboxylase, can entirely reverse the immune recognition phenotype resulting from PIK3CA downregulation. Deleting PCCB in pancreatic tumor cells lacking PIK3CA leads to immune evasion, tumor progression, and ultimately, the death of the host animal. Despite T cells still infiltrating p-αKO tumor sections, there is an enrichment of immune checkpoint molecules such as PD1. Intriguingly, implanting p-αKO cells into PD1-null mice significantly improved mouse survival, hinting at the partial attribution of immune evasion in p-αKO tumors to elevated PD1 expression on T cells. Furthermore, treatment of p-αKO tumors with an anti-PD1 neutralizing antibody yielded comparable results, highlighting the potential applicability of this approach in PDAC treatment, especially when coupled with a PIK3CA inhibitor. Noteworthy outcomes ensued when interfering with the PD-L1/PD1 axis, resulting in T cell-mediated elimination of p-αKO tumor cells. Single-cell sequencing substantiates these findings by confirming that blocking the PD-L1/PD1 interaction reactivates clonal anti-tumor T cells.

Recent investigations underscore the compelling potential of combination therapy involving both PI3K and PD1 inhibitors in cancer treatment. In a study by Isoyama et al. (2020), improved antitumor effects were observed in melanoma and fibrosarcoma models upon the combination of PI3K inhibitors with PD-1 blockade. This therapeutic strategy raised the levels of tumor-specific CD8^+^ T cells thus led to enhanced antitumor activity (44). Similarly, Collins et al. (2021) conducted an *in vivo* expression screen with a pool of known oncogenes and identified PIK3CA as a fasctor promoting resistance to anti-PD1 treatment in a colon carcinoma model. Notably, the resistance to immunotherapy was successfully overturned with the administration of a PI3K inhibitor. These findings offer further support to the potential benefits of combining PI3K inhibitors with PD1 blockade for cancer treatment (45). To further validate the efficacy of this drug combination, future studies can explore its application using our orthotopic implantation model and in genetically modified mouse models that spontaneously develop pancreatic ductal adenocarcinoma (PDAC). This approach ensures a comprehensive understanding of the therapeutic potential in a context that closely mirrors the intricacies of PDAC progression and the TME.

Although the administration of anti-PD1 neutralizing antibody yielded promising results, the treatment alone did not achieve 100% efficacy. Indeed, our scRNA-seq result from PD1-null mice revealed that not all exhausted T cells were reactivated, potentially explaining why some of the mice still succumb to tumor progression despite PD-L1-/PD1 blockade. In addition, to improve survival rates, it is crucial to incorporate other immunotherapies targeting additional checkpoint inhibitory receptors, such as CTLA4, LAG3, and TIGIT. While anti-CTLA4 and anti-PD1 are standard treatments for many solid tumors, TIGIT and LAG3 have emerged as promising new targets for cancer immunotherapy. LAG3, although demonstrating limited efficacy as a monotherapy, is frequently used in combination with other checkpoint inhibitors (11). Similarly, while anti-TIGIT antibodies have shown remarkable effectiveness in treating solid tumors, the dual blockade of PD-1 and TIGIT has demonstrated superior clinical benefits in cancer treatment (48–50). Our IHC data notably revealed a significant upregulation of both TIGIT and LAG3 in p-αKO tumors, suggesting that the inclusion of additional immune checkpoint inhibitors, alongside PD1 inhibition, may further improve treatment outcomes. Therefore, future studies that test blocking other immune checkpoints such as TIGIT, LAG3, CTLA4, or TIM3, in combination with anti-PD1, could provide valuable insights.

While our findings highlight the involvement of PCCB in regulating T cell activities, the precise underlying mechanism remains unclear. Previous studies primarily focused on the role of *Pccb* in the genetic disease propionic acidemia (PA), caused by mutations in *Pccb* or *Pcca* subunits, which impair PCC function (32). Dysfunctional PCC leads to the accumulation of propionyl CoA and other byproducts such as propionylcarnitine (C3), methylcitrate (MC), and 3-hydroxypropionate (3OHPA) (46,47). Intracellular metabolite analysis by LC-MS showed elevated MC and C3 levels upon *Pccb* deletion, although the statistical significance was not reach (Figure 3 – figure supplement 1D). Importantly, intracellular propionyl-CoA levels were similar between αKO and p-αKO cells, potentially due to its conversion to MC and C3. However, the precision of propionyl-CoA detection remains challenging due to its limited quantity. Additional intracellular metabolites assessed through non-targeted LC-MS exhibited noteworthy alterations, suggesting their possible involvement in the regulation of PD1 expression on T cells (Figure 3 – figure supplement 1C). However, further experiments are required to obtain a better understanding of how PCCB-regulated metabolic pathways affect T cell function.

Various studies have highlighted the anti-inflammatory properties of propionate, a byproduct of dysfunctional PCC. Smith et al. (2013) demonstrated that propionic acid regulates colonic Treg (cTreg) homeostasis in the gut lumen. Propionic acid supplementation in the drinking water of germ-free (GF) mice increased cTreg frequency and number in the gut lumen, while Tregs in the spleen, mesenteric lymph nodes, and thymus remained unaffected. Treatment of cTregs isolated from GF mice with propionate led to increased Foxp3 and IL10 expressions, both of which play important roles in Treg-mediated immune suppression. Additionally, cTregs exhibited a higher proliferation rate when co-cultured with propionate (48). Another study by Tedelind et al. (2007) reported that propionate suppressed the secretion of TNFα and IL8 from neutrophils, inhibited the NF-kB pathway, and downregulated immune-related genes in a colon cell line (49). Recently, Hogh et al. reported that propionate upregulated the surface expression of the immune stimulatory NKG2D ligands MHC-class-I-related family (MICA/B) on several cancer cells and activated T cells, mediated by mTORC2 (50). These indirect observations suggest a link between *Pccb* and immune responses although the published results are somewhat conflicting.

Additionally, cytokines and other mechanisms within the TME may contribute to the upregulation of PD1 expression on T cells in p-αKO tumors. RNA sequencing of αKO and p-αKO cells revealed substantial changes in several cytokine-related signaling pathways, indicating potential cytokine involvement in regulating PD1 expression on T cells (Figure 3 - figure supplement 2). Moreover, comparing cells grown *in vitro* to tumor growth *in vivo* poses challenges due to the complex TME and potential crosstalk between immune cells and the metabolic system. Future studies should conduct single-cell RNAseq analysis of *in vivo* tumor cells present within the TME to uncover the mechanisms of immune evasion upon deletion of *Pccb*.

Our study is the first to report an association between PCCB and PIK3CA. We utilized PIK3CA knockout cells (αKO) instead of conventional PIK3CA inhibitors to avoid off-target effects reported in previous studies (51–54), providing a more specific and accurate assessment of PIK3CA’s role in regulating immune responses in PDAC. We employed an orthotopic mouse model for our CRISPR screen, which more closely mimics the TME of human PDAC. Our genome-wide screen study identified PCCB as a key regulator of antitumor T cell activity. We showed that PCCB regulates immune evasion by promoting increased PD-L1 expression in anti-tumor cytotoxic T cells. Consequently, our findings suggest that the resistance of PDAC to PI3K inhibitors and checkpoint inhibitors could potentially be surmounted by deploying both drugs in combination. This may be a promising avenue for future pre-clinical and clinical studies to directly assess if combining these FDA-approved drugs, possibly in conjunction with other treatment modalities, will improve outcomes for pancreatic cancer patients.

## METHODS

### Cell Culture and Reagents

KPC, αKO, and p-αKO were all cultured in complete DMEM media (Gibco, Thermo Fisher Scientific) with 10% FBS (Corning 35-015-CV) and 1% P/S (penicillin/streptomycin: Gibco 15140-122), incubated at 37 °C with 5% CO2. The KPC cell line was a gift from Dr. David Tuveson (Cold Spring Harbor) and was confirmed to possess a G-to-D mutation in the KRAS gene (KRASG12D) and a silent mutation in the transformation related protein 53 gene (TP53R172H) by mutation analysis in our previous study (16). The preparation of KPC and αKO cell lines were descripted before (16). Briefly, KPC cells were infected with lentiviral particles containing CMV-firefly luciferase with a neomycin selection marker (Cellomics Technology, PLV-10064). After 48-hour transfection, KPC cells were selected with 1.5 mg/ml G418, and the luciferase expression was confirmed by the IVIS Lumina III imaging system (Xenogen). KPC cells that express luciferase signals are referred to as WT KPC. *Pik3ca*^-/-^ (αKO) KPC cell lines were generated by transfecting WT KPC cells with *Pik3ca* CRISPR/Cas9 αKO and HDR plasmids (Santa Cruz Biotechnology, sc-422231 and sc-422231-HDR). Transfected cells were selected with 5 mg/mL puromycin and sorted based on red fluorescent protein (RFP) by fluorescence-activated cell sorting (FACS) on a FACSAria (BD Biosciences). RFP^+^ cells were serially diluted in a 96-well plate to generate single cell clones, and clones that showed an absence of PIK3CA protein on western blot were retained (referred to as αKO cell lines). In order to knock out *Pccb* from αKO cell lines, αKO was transfected with *Pccb* CRISPR/Cas9 KO plasmid (Santa Cruz Biotechnology, sc-426258) and incubated for 48 hours. Again, western blot was used to verify PCCB expression and cells without the presence of PCCB and PIK3CA are referred to as p-αKO cell lines. The sgRNA oligonucleotides used for knockout *Pik3ca* are sc-422231 A: GCGCACTATTTATGACCCAG; sc-422231 B: TCACCATGCCGTCATACTCC; sc-422231 C: CAGAAGTCCAAGACTTTCGA. The sgRNA oligonucleotides used for knockout *Pccb* are sc-426258 A: GAGTCATTGAGCCCGATCAC; sc-426258 B: CAGATGTGCCGACTTCGGAA; sc-426258 C: ACTGGACGGGGCCGAATCAA.

### GeCKO v2 Lentiviral Library Preparation and Infection

The mouse GeCKO v2 library was obtained from Addgene (#1000000052). LentiCRISPRv2 is a one-vector plasmid system for the mouse GeCKO (Genome-scale CRISPR Knockout) pooled library. The sublibrary A contains 67,405 sgRNAs targeting 20,611 protein-coding genes, 1175 microRNAs and 1000 control sgRNAs. To ensure no loss of representation, the GeCKO library A was amplified by first electroporating the library to Endura ElectroCompetent cells and then transforming to LB agar plates. The colonies from the LB agar plates were harvested and purified by Maxiprep. To produce lentivirus, the plasmid was used to transfect HEK293FT cells by lipofectamine 2000 plus reagent with packaging plasmids pVSVg and psPAX2. After incubating for 2 days, the lentiviral supernatant from the HEK293FT cells were pooled and filtered to get rid of cellular debris. Then the lentiviral titer was determined through transduction. To calculate the MOI and optimal kill curve, cells were treated with a serial dilution of concentrated virus followed by puromycin selections the next day with titrations of 0, 0.5 mg/ml, 1 mg/ml, and 2 mg/ml. After 48 hours of treatment, cell apoptosis was measured with PI and Hoechst staining to determine the concentrations of viral titer and puromycin that give approximately an MOI of 0.2-0.4 with nearly 100% killing (1:10 dilution). To ensure 350x of library coverage, 1x 10^7^ αKO cells were seeded, and infected with 1:10 dilutions of the viral titer aiming for a MOI of 0.2-0.4.

### CRISPR Knockout Screen in an Orthotopic Pancreatic Implantation Mouse Model

After incubating the transduced αKO cells for 2 days, 7.5x10^5^ of the transfected cells in 30ml of PBS were prepared for each mouse and were implanted in the pancreas of 9 immunocompetent wild-type C57BL/6J mice (B6; Stock#000664). To monitor tumor progression, 100 mg/kg RediJect D-Luciferin (PerkinElmer 770504) was injected intraperitoneally into mice followed by imaging on the IVIS Lumina III imaging system (Xenogen) to monitor tumor progression. Data were analyzed using Living Image v4.3.1 software.

### Next-generation Sequencing and Data Analysis

Genomic DNA was extracted both from the primary tumor and metastatic sites to perform next generation sequencing in order to identify enriched sgRNAs. Qiagen DNeasy Blood & Tissue Kit (#13323) was used to extract DNA according to the manufacturer’s protocol. The concentration of DNA was determined by NanoDrop ND 1000. NGS libraries were prepared by a two-step PCR described before (55,56). All PCR reactions were performed with Herculase II Fusion DNA Polymerase (Agilent K3467). The first PCR (PCR1) utilized primers specific to the sgRNA-expression vector to amplify the sgRNA containing region, in order to preserve full complexity of the GeCKO library. To achieve a 350-fold coverage of the sgRNA library, approximately 200 mg of gDNA for each sample was used, and thus approximate 67 PCR1 reactions were performed for each sample (3 μg gDNA/50 μl reaction). The second PCR (PCR2) is performed to add barcoded adaptors—P7 and P5—to the products from the first PCR, so it allows multiplex sequencing on Illumina NextSeq. For our 67,405 mGeCKOa sgRNA library, at least 7 PCR2 reactions were performed (one 100 μl reaction per 10^4^ constructs in the library). The primers used for 1^st^ round PCR are: PCR1-F: CCCGAGGGGACCCAGAGAG; PCR1-R: GCGCACCGTGGGCTTGTAC. The primers used for the 2^nd^ round PCRs are as follows: Fwd: AATGATACGGCGACCACCGAGATCTACACTCTTTCCCTACACGACGCTCTTCCGATCT-stagger-barcodes-TCTTGTGGAAAGGACGAAACACCG; Rev: CAAGCAGAAGACGGCATAC- GAGAT-barcode-GTGACTGGAGTTCAGACGTGTGCTCTTCCGATCT-stagger-TCTACTATTC- TTTCCCCTGCACTGT. The barcode used for tumor 1 is: AAGTAGAG; and the barcode used for tumor 2 is CGCGCGGT. Before sequencing, DNA quality and quantity were determined by Qubit dsDNA BR Assay Kit and Agilent 2200 TapeStation Analysis. All samples were sequenced on Illumina NextSeq 550 sequencer. The sequencing data was processed for sgRNA representation. The 8bp barcodes were first demultiplexed from the sequencing reads, followed by adapter trimming. The remaining spacer sequences were aligned to the designed sgRNA library by bowtie, with tolerance of a single nucleotide mismatch. The number of mapped sequences were imported into R/RStudio to quantify the total number of reads. Besides, the computational tool MAGeCK (model-based analysis of genome-wide CRISPR-Cas9 knockout) was used to confirm the identified mapping counts.

### Orthotopic Pancreatic Implantation Mouse Model

All the C57BL/6J mice (#000664) and PD-1^-/-^ mice (#028276) were purchased from Jackson Laboratories. The orthotopic implantation surgeries have been described previously (16). Briefly, cells were trypsinized and washed twice with PBS, then counted with a cell counter to prepare 5x10^5^ cells in 30 μl of PBS per mouse for injection (or 7.5 x10^5^ cells for the screen). Mice were anesthetized with a combination of 100 mg/kg ketamine and 10mg/kg xylazine, followed by a small vertical incision made over the left lateral abdominal area. The pancreas was then located with the aid of a light microscope, and the injection was made at the head of pancreas by a sterile Hamilton syringe with a 27-gauge needle. After sutures, mice were given an intraperitoneal injection of 2 mg/kg ketorolac.

### Mouse Survival Studies

Mice implanted with tumor cells (KPC, GeCKO treated αKO, αKO cells, and p-αKO cells) were monitored by IVIS imaging weekly. Mice with weight loss > 15% body weight and/or inability to move were euthanized as these were considered the endpoint. For mice implanted with p-αKO cells, week 8 was set as the endpoint of the experiment and any surviving animals were euthanized.

### Single Cell Isolation

Pancreas with tumor tissues were harvested from mice on day 12 after tumor implantation. After excluding lymph nodes, tissues were washed with cold PBS and minced with scissors in the hood. The minced tissues were washed with PBS, then digested in 5 ml of 1mg/ml Collagenase type V (Worthington LS005282) solution dissolved in HBSS for 20 minutes at 37 °C. 2ml of Roche Red Blood Cell Lysis Buffer (Millipore Sigma 11814389001) was used per sample to get rid of red blood cells, then passed through 40μm cell strainer to generate single cells.

### Enrichment of CD4+ and CD8+ T Cells from Dissociated Tissues

10μl of CD4 (TIL) MicroBeads (Miltenyi Biotec 130-116-475) and 10μl of CD8 (TIL) MicroBeads (Miltenyi Biotec 130-116-478) were added to 90μl of single cell suspensions (up to 10 million cells each) and incubated at 4 °C for 15 minutes. After adding the depletion wash buffer to a final volume of 500μl for up to 50 millions cells, the mixture was separated in a LS column (Miltenyi Biotec 130-042-401) by the magnetic field of the MACS Quadro separator. The flow-through were CD4 and CD8 negative cells, and the magnetically labeled cells that were flushed out after removing the magnetic field were CD4 or CD8 positive T cells. The isolated cells were centrifuged at 300xg 4°C for 5 minutes and resuspended in PBS with 0.1% of BSA to generate 1e6/ml of cells.

### scRNAseq Library Preparation with 10x Genomics Platform

We followed the manufacturer’s protocol provided by 10x Genomics for the preparation of Chromium NextGEM Single Cell 5’ gene expression libraries. Single cell suspensions at a concentration of 0.7-1.2e6/ml were mixed with reverse transcription (RT) reagents, Gel Beads containing barcoded oligonucleotides, and partitioning oil on a microfluidic chip to form Gel Beads in emulsion (GEMs). Within each GEM, a single cell was lysed and the mRNAs barcoded and reverse transcribed to cDNA, followed by PCR amplification of the barcoded cDNAs. Single cell RNA sequencing and VDJ libraries were prepared using the Chromium Next GEM Single Cell 5’ Kit v2 (PN-1000265) and the Chromium Single Cell Mouse TCR Amplification kit (PN-1000254) at the Northport VAMC Single Cell Facility. The gene expression and VDJ libraries were pair-end sequenced (PE150) through a commercial supplier (Novogene, Inc.) at a depth of 20,000 and 5,000 reads/cell, respectively.

### scRNAseq Data Analysis

The Cell Ranger v7.0.0 pipeline was used to demultiplex and align sequencing data to the “ENSEMBL GRCm39” mouse transcriptome to generate gene expression matrices. The Cell Ranger VDJ pipeline was used to assemble sequences and identify paired clonotypes on the V(D)J libraries. The gene expression matrices were further analyzed by the Seurat R package. Three quality control criteria were employed to filter the matrices to exclude unwanted sources of variations: number of detected transcripts, genes, and percent of reads mapping to mitochondrial genes. Cells with UMIs over the interquartile range of 95% (potential doublets) and under 1000 (potential fragments) or a mitochondrial proportion higher than 20% (potential apoptotic) were removed. Moreover, we used the Doublet Finder R algorithm to further eliminate doublet contamination. After the quality control, we annotated cellular identity using the R package SingleR, which assigns each cell to a reference type that has the most similar expression profile with. In further analysis, sctransform package in Seurat was used to normalize UMI count data. The principal component analysis (PCA) of high-dimensional data was performed to identify highly variable genes in each sample, and the top principal components were selected for unsupervised clustering of cells with a graph-based clustering and visualized with Uniform Manifold Approximation and Projection (UMAP).

### Western Blotting

Cells were washed twice by PBS followed by lysis in RIPA buffer (50 mM HEPES, pH 7.4, 10 mM sodium pyrophosphate, 50 mM NaF, 5 mM EDTA, 1 mM sodium orthovanadate, 0.25% sodium deoxycholate, 1% NP40, 1 mM PMSF, and protease inhibitor cocktail (Sigma P8340). After centrifuge, cell lysates were collected, and proteins were separated by SDS-PAGE gel followed by semi-dry transfer onto nitrocellulose membranes. After blocking with 5% milk in tris-buffered saline plus 0.1% Tween 20 (TBST), primary antibodies prepared in a 1:1000 dilution with TBST for overnight. After incubating the membrane with horseradish peroxidase (HRP) linked secondary antibodies (Thermo Fisher Scientific #62-6520 & #31460) in a 1:5000 dilution with TBST, signals were developed by adding ECL reagent or SuperSignal West Femto (Thermo Scientific 34095). Signals were detected by a FluorChem E imager (ProteinSimple). Primary antibodies used: PCCB: Millipore sigma HPA036939; PIK3CA: Cell Signaling #4249S; HSP90: Cell Signaling #4874S; PTEN: Cell Signaling #9559S; pAKT (Thr308): Cell Signaling #9275S; pAKT (Ser473): Cell Signaling #4051S; AKT 1/2/3: Santa Cruz Biotechnology sc-377556; actin: Millipore Sigma #A2103; PPRAS40: Cell Signaling #2997S.

### Cell Proliferation Assay

2500 cells were plated in triplicates for each cell line on a 12-well plate on day 0. From day 1 to day 4, cells were washed with PBS, trypsinized and resuspended with completed media and then stained with trypan blue (Invitrogen T10282) with a 1:1 dilution. The cells were then counted using Countess II automated cell counter (Life Technologies). Cell numbers were plotted by GraphPad Prism 9.

### Real-time Metabolic Analysis

The Seahorse XF96 extracellular flux analyzer (Seahorse Bioscience, USA) was utilized to measure the oxygen consumption rate (OCR). Cells were cultured in 96-well XF microplates with 20,000 cells per well and incubated for 18 hours at 37°C. To evaluate mitochondrial activity, sequential injections of oligomycin, the electron transport chain uncoupler FCCP, and specific inhibitor of the mitochondrial respiratory chain were performed. The OCR data were normalized to cell number.

### LC-MS

To assess the total metabolite levels, cells were plated in 6-well plates in triplicates with equal cell numbers for each well. After cells were washed three times with PBS, a mixture of methanol, acetonitrile, and water with formic acid was quickly added to the dishes. The plates were then placed on ice for 5 minutes, followed by the addition of 15% NH_4_HCO_3_ to neutralize the acetic acid. The cells were collected by scraping the centrifuged, and the resulting supernatant was used for LC-MS analysis. The LC-MS employed hydrophilic interaction chromatography (HILIC) coupled with electrospray ionization to a Q Exactive PLUS hybrid quadrupole orbitrap mass spectrometer (Thermo Scientific). A specific LC separation was carried out using an XBridge BEH Amide column, with a gradient of solvent A and solvent B. The gradient varied over a specific time frame, and the flow rate, injection volume, and column temperature were controlled. The metabolite features were extracted using MAVEN software, considering the labeled isotope and a mass accuracy window. The isotope natural abundance and tracer isotopic impurity were corrected using AccuCor. The LC-MS and data analysis was done by Metabolomics Shared Resource at the Rutgers Cancer Institute of New Jersey.

### Histology

Mouse tumors were fixed in 10% formalin for 24 hours. Tissue processing was done by Leica ASP300S Tissue Processor. After embedding with paraffin, tissues were sectioned into 4μm sections. H&E staining was performed using Hematoxylin and eosin.

Immunohistochemistry (IHC) was performed by deparaffinization and rehydration of the tissue sections, followed by antigen retrieval in citrate buffer, pH 6.0 (Vector Laboratories H-3300-250) using a Decloaking Chamber (Biocare Medical). To block endogenous peroxidase activity, 3% H_2_O_2_ (Fisher Scientific 7722-84-1) was used. And endogenous biotin was blocked by using the Avidin/Biotin Blocking kit (Vector Laboratories SP-2001). After incubation of primary antibodies (1:1000) overnight, the signal was developed by the R.T.U. Vectastain Kit (Vector PK-7100) and 3,3’-diaminobenzidine in chromogen solution (Agilent K3467). Hematoxylin (Agilent CS70030-2) then used to counterstain the nucleus. Antibodies used in IHC were CD3: Abcam #ab16669; CD4: Abcam #ab183685; CD8α: Cell Signaling #98941S; PD1: Abcam #ab214421; CTLA4: Abcam #ab237712; LAG3: Abcam #ab209238; TIGIT: Abcam #ab300073; TIM3: Abcam #ab2413332.

### IHC Quantification

Two consecutive sections were obtained from each sample. Each section two or three images with 40x magnification were taken on an Olympus BX43 with DP26 camera. The number of cells were manually counted by QuPath.

### Anti-PD1 Antibody Treatment

The InVivoPlus anti-mouse PD1 (CD279) antibody was purchased from BioxCell (#BP0273). The PD1 antibody was diluted in InVivoPure pH7 Dilution buffer (BioxCell #IP0070) to make a 12.5 μg/mg (average of 22 mg mice) solution. After mice were implanted with p-αKO cells, 100μl of prepared mixture or the dilution buffer were injected intraperitoneally to mice weekly for 8 weeks. Mice were imaged by IVIS from week 5 to week 8, and mice survival data were recorded for survival studies.

### Statistical Analysis

All statistical analyses were done by Graphpad Prism 9. For all statistical comparisons, 2-tailed Student’s *t* test was used for comparison between 2 groups. The one-way ANOVA with Bonferroni’s post hoc test and Kruskal-Wallis test with Dunn’s post hoc test were used for multi-group comparisons. *p* values less than 0.05 were considered statistically significant.

### Study Approval

All experiments in mice were conducted in accordance with the Office of Laboratory Animal Welfare and approved by the IACUC of Stony Brook University, Stony Brook, New York.

### Data Availability

scRNAseq data from figure 1 and 6 have been deposited in GEO under accession code GSE254041. CRISPR Screen data from figure 2 have been deposited in BioProject under the accession code PRJNA1068774. The data analysis computer code for scRNAseq is available upon request.

## ACKNOWLEDGMENTS

We express our sincere appreciation to our dedicated laboratory scientist, Lisa Ballou and Giuseppe Caso, as well as our undergraduate research assistant Mark Koch and Nathaniel Tchangou for their unwavering commitment, expertise, and contributions to this project.

## ARTICLE AND AUTHOR INFORMATION

### Han Victoria Han

Department of Physiology and Biophysics, Stony Brook University, Stony Brook, New York Department of Biomedical Engineering, Stony Brook University, Stony Brook, New York **Contribution:** Conceptualization, Methodology, Software, Validation, Formal Analysis, Investigation, Writing-Original Draft Preparation, Writing-Review & Editing, Data Curation, Visualization

**Competing interests:** No competing interests declared

### Richard Efem

Northport Veteran Affair Medical Center, Northport, New York, USA

**Contribution:** Software, Visualization

**Competing interests:** No competing interests declared

### Barbara Rosati

Department of Physiology and Biophysics, Stony Brook University, Stony Brook, New York Northport Veteran Affair Medical Center, Northport, New York, USA

**Contribution:** Methodology, Writing-Review & Editing

**Competing interests:** No competing interests declared

### Kevin Lu

Department of Chemical Biology, Ernest Mario School of Pharmacy, Rutgers-The State University of New Jersey, Piscataway, New Jersey

**Contribution:** Methodology, Resources

**Competing interests:** No competing interests declared

### Sara Maimouni

Department of Chemical Biology, Ernest Mario School of Pharmacy, Rutgers-The State University of New Jersey, Piscataway, New Jersey

**Contribution:** Methodology, Resources

**Competing interests:** No competing interests declared

### Ya-Ping Jiang

Department of Physiology and Biophysics, Stony Brook University, Stony Brook, New York

**Contribution:** Methodology

**Competing interests:** No competing interests declared

### Valeria Montoya

Department of Microbiology and Immunology, Renaissance School of Medicine at Stony Brook University, Stony Brook, New York, USA

Center for Infectious Diseases, Renaissance School of Medicine at Stony Brook University, Stony Brook, New York, USA

**Contribution:** Methodology, Resources

**Competing interests:** No competing interests declared

### Adrianus W. M. Van Der Velden

Department of Microbiology and Immunology, Renaissance School of Medicine at Stony Brook University, Stony Brook, New York, USA

Center for Infectious Diseases, Renaissance School of Medicine at Stony Brook University, Stony Brook, New York, USA

**Contribution:** Methodology, Resources

**Competing interests:** No competing interests declared

### Wei-Xing Zong

Department of Chemical Biology, Ernest Mario School of Pharmacy, Rutgers-The State University of New Jersey, Piscataway, New Jersey

**Contribution:** Methodology, Resources, Writing-Review & Editing

**Competing interests:** No competing interests declared

### Richard Z. Lin

Department of Physiology and Biophysics, Stony Brook University, Stony Brook, New York Northport Veteran Affair Medical Center, Northport, New York, USA

**Contribution:** Conceptualization, Funding Acquisition, Project Administration, Supervision, Writing-Review & Editing

**Competing interests:** No competing interests declared

### Impact Statement

Genome-wide CRISPR screen successfully identified propionyl-CoA subunit B as a key regulator for anti-tumor T cell activity in pancreatic cancer, which holds promise for advancing immunotherapy strategies in curing pancreatic cancer.

## Funding Information

This study was supported in part by Department of Veterans Affairs Merit Review BX004083 (R.Z.L.) and National Institute of Health R21CA274425 (R.Z.L.), R01CA129536 (W.X.Z.) and R01CA236242 (W.X.Z.).

The funders had no role in study design, data collection and interpretation, or the decision to submit the work for publication.

---

**Figure 3 – figure supplement 1.**
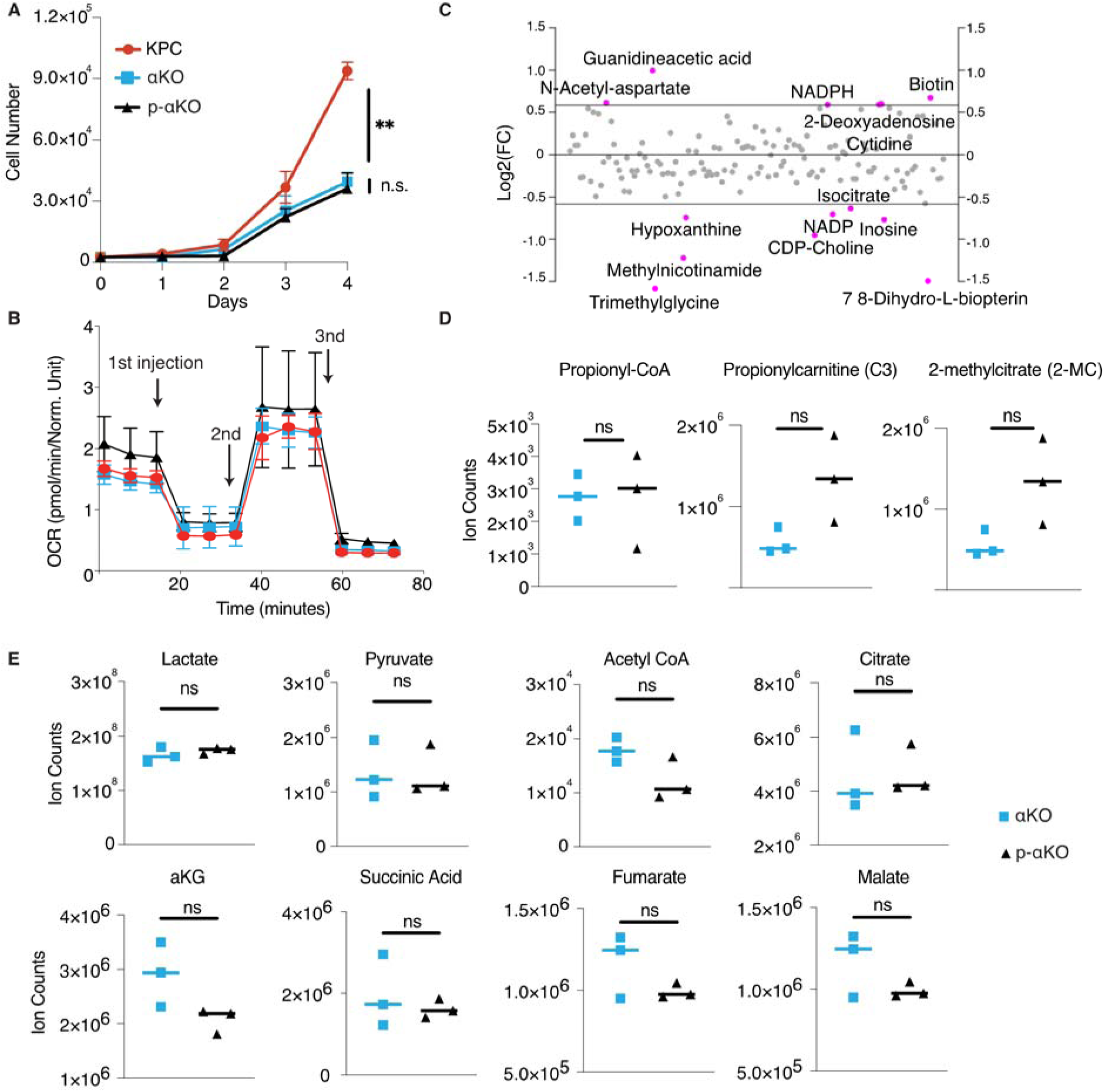
*In vitro* characterization of p-αKO cells. **(A)** Growth rates of KPC, αKO and p-αKO cell lines in culture were measured from day 1 to day 4 (mean ± SD, n=3). ***P= 0.0009 for KPC vs. p-αKO; not significant (ns) for αKO vs. p-αKO (One-way ANOVA). **(B)** Mitochondria function was evaluated by measuring the oxygen consumption rate (OCR) with Seahorse. 1st injection: oligomycin; 2nd injection: FCCP; 3rd injection: antimycin A & rotenone. **(C)** Metabolic changes between p-αKO and αKO revealed by non-targeted LC-MS analysis. Each data point represents the Log2 fold change of p-αKO over αKO, with a threshold set at 1.5-fold changes. Metabolites with differences exceeding this threshold were depicted in red. **(D)** LC-MS exploring changes of propionyl-CoA, propionylcarnitine, and 2-methylcitrate. Data are represented as ion counts ± SD (N=3 for all groups). ns: not significant. **(E)** LC-MS exploring changes of metabolites of the TCA cycle. Data are represented as ion counts ± SD (N=3 for all groups). ns: not significant.

**Figure 4 – figure supplement 1.**
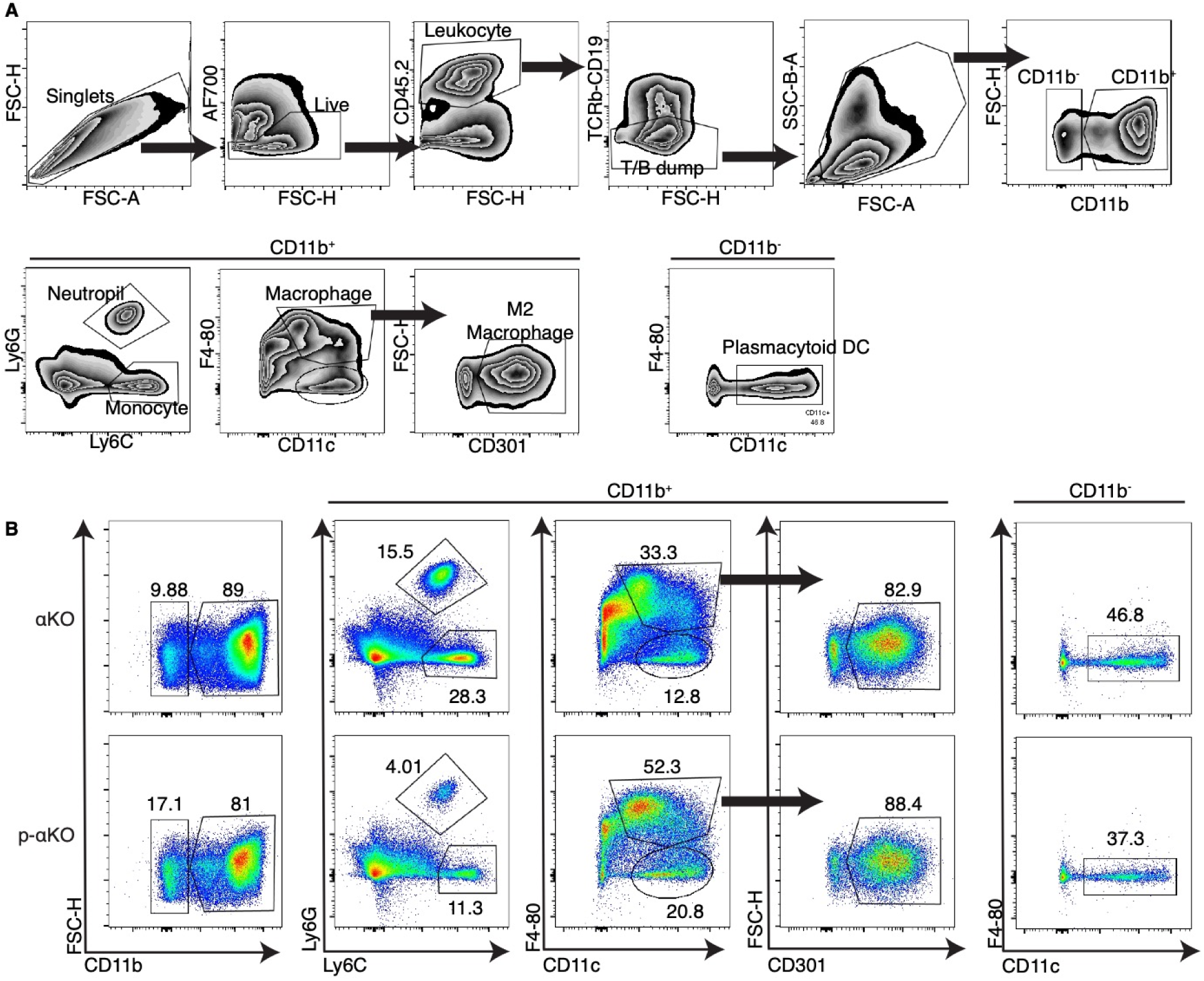
Flow cytometry of myeloid cells infiltrated in αKO vs. p-αKO. **(A)** The gating strategy of the myeloid panel. **(B)** Flow cytometry analysis of the cells isolated from αKO and p-αKO tumors implanted in C57BL/6J mice. The numbers presented in the figures represent cell percentages relative to the total cell counts from the respective previous groups.

**Figure 3 - figure supplement 2.**
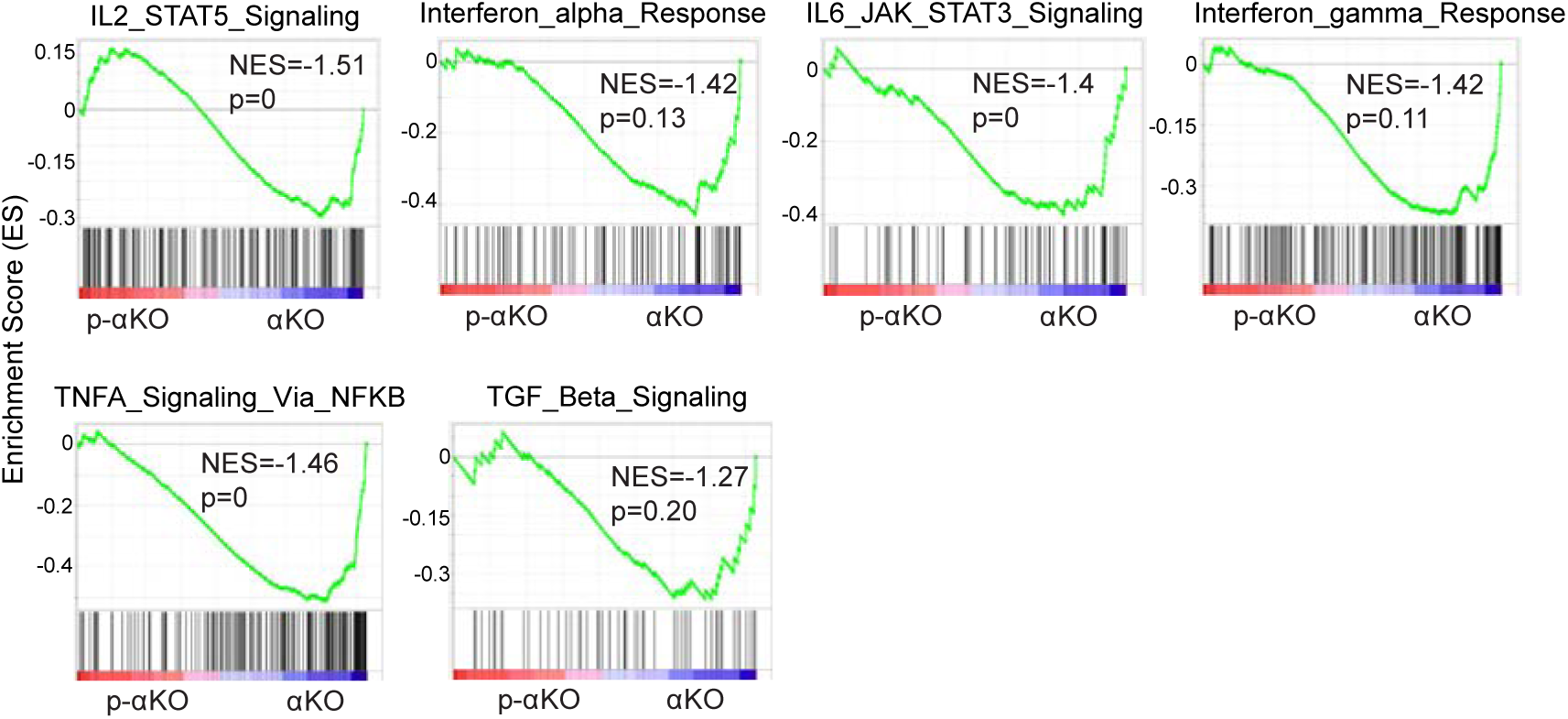
Gene set enrichment analysis (GSEA) of cytokine-related pathways in αKO cells (n=3) vs. p-αKO cells (n=3). These pathways include IL2-STAT5 signaling pathway, interferon α response, IL6-JAK-STAT3 signaling pathway, interferon γ response, TNFA signaling via NFKB, and TGF β signaling pathway. NES: normalized enrichment score.

## Notes

### Competing Interest Statement

The authors have declared no competing interest.

### Summary of Updates

Major revision to the manuscript with additional experiments. New authors also added

## REFERENCES

1. Siegel RL, Miller KD, Wagle NS, Jemal A. Cancer statistics, 2023. CA Cancer J Clin. Jan 2023;73(1):17–48. doi:10.3322/caac.21763

2. Bian J, Almhanna K. Pancreatic cancer and immune checkpoint inhibitors-still a long way to go. Transl Gastroenterol Hepatol. 2021;6:6. doi:10.21037/tgh.2020.04.03

3. Mucileanu A, Chira R, Mircea PA. PD-1/PD-L1 expression in pancreatic cancer and its implication in novel therapies. Med Pharm Rep. Oct 2021;94(4):402–410. doi:10.15386/mpr-2116

4. Balsano R, Zanuso V, Pirozzi A, Rimassa L, Bozzarelli S. Pancreatic Ductal Adenocarcinoma and Immune Checkpoint Inhibitors: The Gray Curtain of Immunotherapy and Spikes of Lights. Curr Oncol. Mar 30 2023;30(4):3871–3885. doi:10.3390/curroncol30040293

5. Fan J-q, Wang M-F, Chen H-L, Shang D, Das JK, Song J. Current advances and outlooks in immunotherapy for pancreatic ductal adenocarcinoma. Molecular Cancer. 2020/02/15 2020;19(1):32. doi:10.1186/s12943-020-01151-3

6. Johnson BA, Yarchoan M, Lee V, Laheru DA, Jaffee EM. Strategies for Increasing Pancreatic Tumor Immunogenicity. Clinical Cancer Research. 2017;23(7):1656–1669. doi:10.1158/1078-0432.ccr-16-2318

7. Halbrook CJ, Lyssiotis CA, Pasca di Magliano M, Maitra A. Pancreatic cancer: Advances and challenges. Cell. Apr 13 2023;186(8):1729–1754. doi:10.1016/j.cell.2023.02.014

8. Liu L, Huang X, Shi F, et al. Combination therapy for pancreatic cancer: anti-PD-(L)1- based strategy. J Exp Clin Cancer Res. Feb 9 2022;41(1):56. doi:10.1186/s13046-022-02273-w

9. Li H-B, Yang Z-H, Guo Q-Q. Immune checkpoint inhibition for pancreatic ductal adenocarcinoma: limitations and prospects: a systematic review. Cell Communication and Signaling. 2021/11/24 2021;19(1):117. doi:10.1186/s12964-021-00789-w

10. Freed-Pastor WA, Lambert LJ, Ely ZA, et al. The CD155/TIGIT axis promotes and maintains immune evasion in neoantigen-expressing pancreatic cancer. Cancer Cell. Oct 11 2021;39(10):1342–1360.e14. doi:10.1016/j.ccell.2021.07.007

11. Gulhati P, Schalck A, Jiang S, et al. Targeting T cell checkpoints 41BB and LAG3 and myeloid cell CXCR1/CXCR2 results in antitumor immunity and durable response in pancreatic cancer. Nature Cancer. 2023/01/01 2023;4(1):62–80. doi:10.1038/s43018-022-00500-z

12. Panni RZ, Herndon JM, Zuo C, et al. Agonism of CD11b reprograms innate immunity to sensitize pancreatic cancer to immunotherapies. Sci Transl Med. Jul 3 2019;11(499)doi:10.1126/scitranslmed.aau9240

13. Vonderheide RH. CD40 Agonist Antibodies in Cancer Immunotherapy. Annu Rev Med. Jan 27 2020;71:47–58. doi:10.1146/annurev-med-062518-045435

14. Zhu Y, Herndon JM, Sojka DK, et al. Tissue-Resident Macrophages in Pancreatic Ductal Adenocarcinoma Originate from Embryonic Hematopoiesis and Promote Tumor Progression. Immunity. Aug 15 2017;47(2):323–338.e6. doi:10.1016/j.immuni.2017.07.014

15. Wu Y, Yi M, Niu M, Mei Q, Wu K. Myeloid-derived suppressor cells: an emerging target for anticancer immunotherapy. Mol Cancer. Sep 26 2022;21(1):184. doi:10.1186/s12943-022-01657-y

16. Sivaram N, McLaughlin PA, Han HV, et al. Tumor-intrinsic PIK3CA represses tumor immunogenicity in a model of pancreatic cancer. J Clin Invest. May 21 2019;129(8):3264–3276. doi:10.1172/jci123540

17. Baleeiro RB, Bouwens CJ, Liu P, et al. MHC class II molecules on pancreatic cancer cells indicate a potential for neo-antigen-based immunotherapy. Oncoimmunology. 2022;11(1):2080329. doi:10.1080/2162402x.2022.2080329

18. Yamamoto K, Venida A, Yano J, et al. Autophagy promotes immune evasion of pancreatic cancer by degrading MHC-I. Nature. 2020/05/01 2020;581(7806):100–105. doi:10.1038/s41586-020-2229-5

19. Hegde S, Krisnawan VE, Herzog BH, et al. Dendritic Cell Paucity Leads to Dysfunctional Immune Surveillance in Pancreatic Cancer. Cancer Cell. Mar 16 2020;37(3):289–307.e9. doi:10.1016/j.ccell.2020.02.008

20. Jiang H, Hegde S, Knolhoff BL, et al. Targeting focal adhesion kinase renders pancreatic cancers responsive to checkpoint immunotherapy. Nat Med. Aug 2016;22(8):851–60. doi:10.1038/nm.4123

21. Özdemir BC, Pentcheva-Hoang T, Carstens JL, et al. Depletion of carcinoma-associated fibroblasts and fibrosis induces immunosuppression and accelerates pancreas cancer with reduced survival. Cancer Cell. Jun 16 2014;25(6):719–34. doi:10.1016/j.ccr.2014.04.005

22. Shi C, Washington MK, Chaturvedi R, et al. Fibrogenesis in pancreatic cancer is a dynamic process regulated by macrophage-stellate cell interaction. Lab Invest. Apr 2014;94(4):409–21. doi:10.1038/labinvest.2014.10

23. Johnson BA, 3rd, Yarchoan M, Lee V, Laheru DA, Jaffee EM. Strategies for Increasing Pancreatic Tumor Immunogenicity. Clin Cancer Res. Apr 1 2017;23(7):1656–1669. doi:10.1158/1078-0432.Ccr-16-2318

24. Popovic A, Jaffee EM, Zaidi N. Emerging strategies for combination checkpoint modulators in cancer immunotherapy. J Clin Invest. Aug 1 2018;128(8):3209–3218. doi:10.1172/jci120775

25. Mace TA, Shakya R, Pitarresi JR, et al. IL-6 and PD-L1 antibody blockade combination therapy reduces tumour progression in murine models of pancreatic cancer. Gut. Feb 2018;67(2):320–332. doi:10.1136/gutjnl-2016-311585

26. Dey P, Li J, Zhang J, et al. Oncogenic KRAS-Driven Metabolic Reprogramming in Pancreatic Cancer Cells Utilizes Cytokines from the Tumor Microenvironment. Cancer Discov. Apr 2020;10(4):608–625. doi:10.1158/2159-8290.Cd-19-0297

27. Balachandran VP, Łuksza M, Zhao JN, et al. Identification of unique neoantigen qualities in long-term survivors of pancreatic cancer. Nature. Nov 23 2017;551(7681):512–516. doi:10.1038/nature24462

28. Moore AR, Rosenberg SC, McCormick F, Malek S. RAS-targeted therapies: is the undruggable drugged? Nat Rev Drug Discov. Aug 2020;19(8):533–552. doi:10.1038/s41573-020-0068-6

29. Wu CY, Carpenter ES, Takeuchi KK, et al. PI3K regulation of RAC1 is required for KRAS-induced pancreatic tumorigenesis in mice. Gastroenterology. Dec 2014;147(6):1405–16.e7. doi:10.1053/j.gastro.2014.08.032

30. Tu AA, Gierahn TM, Monian B, et al. TCR sequencing paired with massively parallel 3′ RNA-seq reveals clonotypic T cell signatures. Nature Immunology. 2019;20(12):1692–1699. doi:10.1038/s41590-019-0544-5

31. Andreatta M, Corria-Osorio J, Müller S, Cubas R, Coukos G, Carmona SJ. Interpretation of T cell states from single-cell transcriptomics data using reference atlases. Nature Communications. 2021/05/20 2021;12(1):2965. doi:10.1038/s41467-021-23324-4

32. Wongkittichote P, Ah Mew N, Chapman KA. Propionyl-CoA carboxylase - A review. Mol Genet Metab. Dec 2017;122(4):145–152. doi:10.1016/j.ymgme.2017.10.002

33. Shchelochkov OA, Carrillo N, Venditti C. Propionic Acidemia. In: Adam MP, Everman DB, Mirzaa GM, et al, eds. GeneReviews(®). University of Washington, Seattle Copyright © 1993-2022, University of Washington, Seattle. GeneReviews is a registered trademark of the University of Washington, Seattle. All rights reserved.; 1993.

34. Grünert SC, Müllerleile S, De Silva L, et al. Propionic acidemia: clinical course and outcome in 55 pediatric and adolescent patients. Orphanet J Rare Dis. Jan 10 2013;8:6. doi:10.1186/1750-1172-8-6

35. Kallies A, Zehn D, Utzschneider DT. Precursor exhausted T cells: key to successful immunotherapy? Nat Rev Immunol. Feb 2020;20(2):128–136. doi:10.1038/s41577-019-0223-7

36. Im SJ, Hashimoto M, Gerner MY, et al. Defining CD8+ T cells that provide the proliferative burst after PD-1 therapy. Nature. Sep 15 2016;537(7620):417–421. doi:10.1038/nature19330

37. Utzschneider DT, Charmoy M, Chennupati V, et al. T Cell Factor 1-Expressing Memory-like CD8(+) T Cells Sustain the Immune Response to Chronic Viral Infections. Immunity. Aug 16 2016;45(2):415–27. doi:10.1016/j.immuni.2016.07.021

38. Sun P, Zhang X, Wang R-J, et al. PI3Kα inhibitor CYH33 triggers antitumor immunity in murine breast cancer by activating CD8+T cells and promoting fatty acid metabolism. Journal for ImmunoTherapy of Cancer. 2021;9(8):e003093. doi:10.1136/jitc-2021-003093

39. Peng W, Chen JQ, Liu C, et al. Loss of PTEN Promotes Resistance to T Cell-Mediated Immunotherapy. Cancer Discov. Feb 2016;6(2):202–16. doi:10.1158/2159-8290.Cd-15-0283

40. Liu Y-T, Sun Z-J. Turning cold tumors into hot tumors by improving T-cell infiltration. Theranostics. 2021;11(11):5365–5386. doi:10.7150/thno.58390

41. Borcoman E, De La Rochere P, Richer W, et al. Inhibition of PI3K pathway increases immune infiltrate in muscle-invasive bladder cancer. Oncoimmunology. 2019;8(5):e1581556. doi:10.1080/2162402x.2019.1581556

42. Sai J, Owens P, Novitskiy SV, et al. PI3K Inhibition Reduces Mammary Tumor Growth and Facilitates Antitumor Immunity and Anti-PD1 Responses. Clin Cancer Res. Jul 1 2017;23(13):3371–3384. doi:10.1158/1078-0432.Ccr-16-2142

43. Sun P, Meng L-H. Emerging roles of class I PI3K inhibitors in modulating tumor microenvironment and immunity. Acta Pharmacologica Sinica. 2020;41(11):1395–1402. doi:10.1038/s41401-020-00500-8

44. Isoyama S, Mori S, Sugiyama D, et al. Cancer immunotherapy with PI3K and PD-1 dual-blockade via optimal modulation of T cell activation signal. J Immunother Cancer. Aug 2021;9(8)doi:10.1136/jitc-2020-002279

45. Collins NB, Al Abosy R, Miller BC, et al. PI3K activation allows immune evasion by promoting an inhibitory myeloid tumor microenvironment. Journal for ImmunoTherapy of Cancer. 2022;10(3):e003402. doi:10.1136/jitc-2021-003402

46. Kurczynski TW, Hoppel CL, Goldblatt PJ, Gunning WT. Metabolic studies of carnitine in a child with propionic acidemia. Pediatr Res. Jul 1989;26(1):63–6. doi:10.1203/00006450-198907000-00018

47. Ando T, Rasmussen K, Wright JM, Nyhan WL. Isolation and identification of methylcitrate, a major metabolic product of propionate in patients with propionic acidemia. J Biol Chem. Apr 10 1972;247(7):2200–4.

48. Smith PM, Howitt MR, Panikov N, et al. The microbial metabolites, short-chain fatty acids, regulate colonic Treg cell homeostasis. Science. Aug 2 2013;341(6145):569–73. doi:10.1126/science.1241165

49. Tedelind S, Westberg F, Kjerrulf M, Vidal A. Anti-inflammatory properties of the short-chain fatty acids acetate and propionate: a study with relevance to inflammatory bowel disease. World J Gastroenterol. May 28 2007;13(20):2826–32. doi:10.3748/wjg.v13.i20.2826

50. Høgh RI, Møller SH, Jepsen SD, et al. Metabolism of short-chain fatty acid propionate induces surface expression of NKG2D ligands on cancer cells. The FASEB Journal. 2020;34(11):15531–15546. doi:10.1096/fj.202000162r

51. Wang Y, Kuramitsu Y, Baron B, et al. PI3K inhibitor LY294002, as opposed to wortmannin, enhances AKT phosphorylation in gemcitabine-resistant pancreatic cancer cells. Int J Oncol. Feb 2017;50(2):606–612. doi:10.3892/ijo.2016.3804

52. Gharbi SI, Zvelebil MJ, Shuttleworth SJ, et al. Exploring the specificity of the PI3K family inhibitor LY294002. Biochem J. May 15 2007;404(1):15–21. doi:10.1042/bj20061489

53. Miller MS, Thompson PE, Gabelli SB. Structural Determinants of Isoform Selectivity in PI3K Inhibitors. Biomolecules. Feb 26 2019;9(3)doi:10.3390/biom9030082

54. Kong D, Yamori T. Phosphatidylinositol 3-kinase inhibitors: promising drug candidates for cancer therapy. Cancer Sci. Sep 2008;99(9):1734–40. doi:10.1111/j.1349-7006.2008.00891.x

55. Chen S, Sanjana NE, Zheng K, et al. Genome-wide CRISPR screen in a mouse model of tumor growth and metastasis. Cell. Mar 12 2015;160(6):1246–60. doi:10.1016/j.cell.2015.02.038

56. Joung J, Konermann S, Gootenberg JS, et al. Genome-scale CRISPR-Cas9 knockout and transcriptional activation screening. Nature Protocols. 2017;12(4):828–863. doi:10.1038/nprot.2017.016

